# Genomic Analysis of *Blomia tropicalis* Identifies Novel Allergens for Component-Resolved Diagnosis of Mite Allergy

**DOI:** 10.1101/2023.02.09.527948

**Authors:** Qing Xiong, Xiaoyu Liu, Angel Tsz-Yau Wan, Nat Malainual, Xiaojun Xiao, Hui Cao, Man-Fung Tang, Judy Kin-Wing Ng, Soo-Kyung Shin, Yang Yie Sio, Mingqiang Wang, Baoqing Sun, Ting-Fan Leung, Fook Tim Chew, Anchalee Tungtrongchitr, Stephen Kwok-Wing Tsui

## Abstract

**Background:** *Blomia* (*B.*) *tropicalis*, as an important species of house dust mites (HDMs), plays a critical role in allergic diseases in tropical populations, but its allergen components are less investigated than those of other HDMs. Multiple omics methods have largely improved the identification of mite allergens. Here, we sought to identify a comprehensive allergen profile of *B. tropicalis* and advance the allergen component-resolved diagnosis (CRD) of mite allergy.

**Methods:** Reference mite allergen sequences were searched in a high-quality genome of *B. tropicalis*. Comparative analysis was performed for important allergen groups. ELISA was used to assess the allergenicities of recombinant proteins of specific allergens.

**Results:** A complete allergen profile of *B. tropicalis* was revealed, including thirty-seven allergen groups (up to Blo t 42). In-depth comparative analysis not only determined the homology of major allergen groups 5 and 21 but also shed light on the emergence and divergence of chitin-binding allergens. The specific Blo t 12 was identified to be a chitin-binding protein originating from the chitinase of allergen group 15. Immunoassays of recombinant proteins verified three novel allergens and the ELISA results suggested geographical differences in the *B. tropicalis* sensitization rate.

**Conclusions:** The comprehensive allergen profile revealed in *B. tropicalis*, the comparative analysis of allergen groups and the immunoassay assessment of recombinant proteins largely expanded our knowledge to *B. tropicalis* allergens and could ultimately benefit the CRD of HDM allergy.

## Introduction

House dust mites (HDMs) are one of the major causes of human allergic diseases, including asthma, allergic rhinitis, and atopic dermatitis. The prevalence of HDM allergies highly varies among different climates, urbanization rates and hygiene situations (1). It was reported that the overall sensitization rate of HDMs in Asian countries can be up to 90% in atopic patients and was higher than that in Western countries (2). *B. tropicalis* was considered a storage mite but is now identified as an important species of HDM and causes a series of allergic diseases, especially in tropical regions such as Singapore (3). Compared with the major HDM species *Dermatophagoides* (*D.*) *farinae* and *D. pteronyssinus*, *B. tropicalis* has much fewer reported allergen groups (4–6). Because of the increasing sensitization rates to *B. tropicalis* over the last decades (7–10), comprehensive identification of its allergen components has become an urgent task when component-resolved diagnosis (CRD) has become an inevitable trend (5, 11).

In this study, genome-wide analysis identified a comprehensive putative allergen profile of *B. tropicalis*, including many novel groups, and cross-species comparative analysis revealed a range of allergen gene family variations. Moreover, three novel allergens were identified by ELISA experiments using serum samples from patients with sensitization to *B. tropicalis*. Collectively, this genome-wide analysis of *B. tropicalis* revealed a global picture of the allergen profile and provided insights into the CRD of mite allergy.

## Methods

### Genome and transcriptome data

The genome and transcriptome data of *B. tropicalis* have been reported (11) and deposited under in the NCBI database under BioProject accession PRJNA702011. The GenBank assembly accession is GCA_021650745.2 and the SRA accessions of transcriptome data are SRR13742047, SRR13742048, SRR13742049 and SRR13742050.

### In silico identification of allergens

The protein sequences of allergens in astigmatic mites were downloaded from the WHO/IUIS allergen nomenclature database (4) (dated March 2022) as references. The reference allergens were searched in the annotated protein sequences of *B. tropicalis* genome (11) using BLASTP v2.9.0 (12) with the options ‘-evalue 1e-6 -max_hsps 1 -max_target_seqs 1 -outfmt 6’. The top matched proteins were considered as candidate allergens. Then, the transcriptome data were mapped to the genome using Hisat2 (13) and the format conversion was performed by SAMtools (14). The sorted and indexed bam file was visualized in the Integrative Genomics Viewer (IGV) (15) and used for the manual curation of the candidate allergens based on transcriptome data to confirm intron-exon split sites. The final *in silico* identified allergens were listed in Table 1 and the sequences have been deposited in NCBI GenBank database (accessions are OQ285922–OQ285989).

**Table 1.**
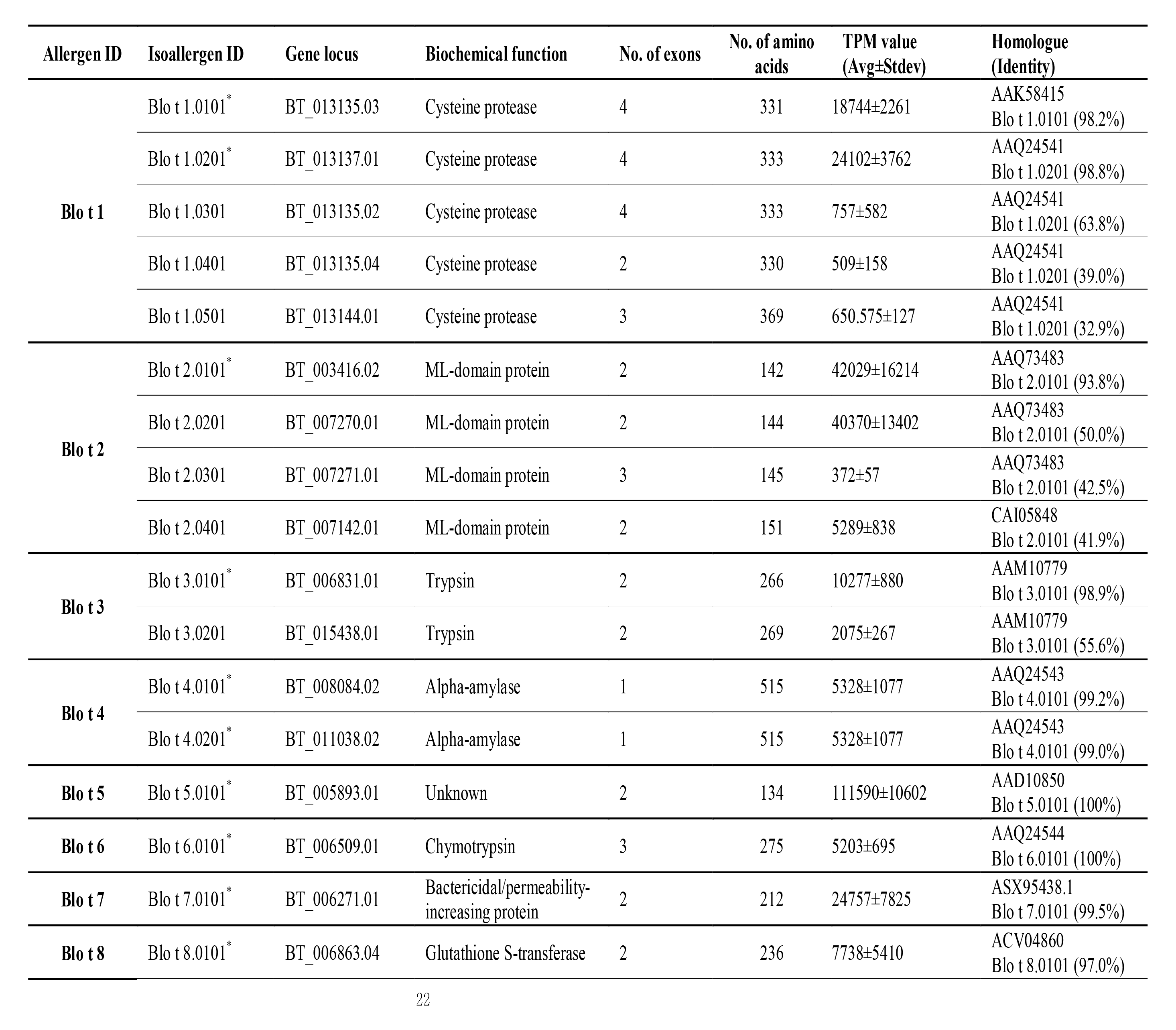
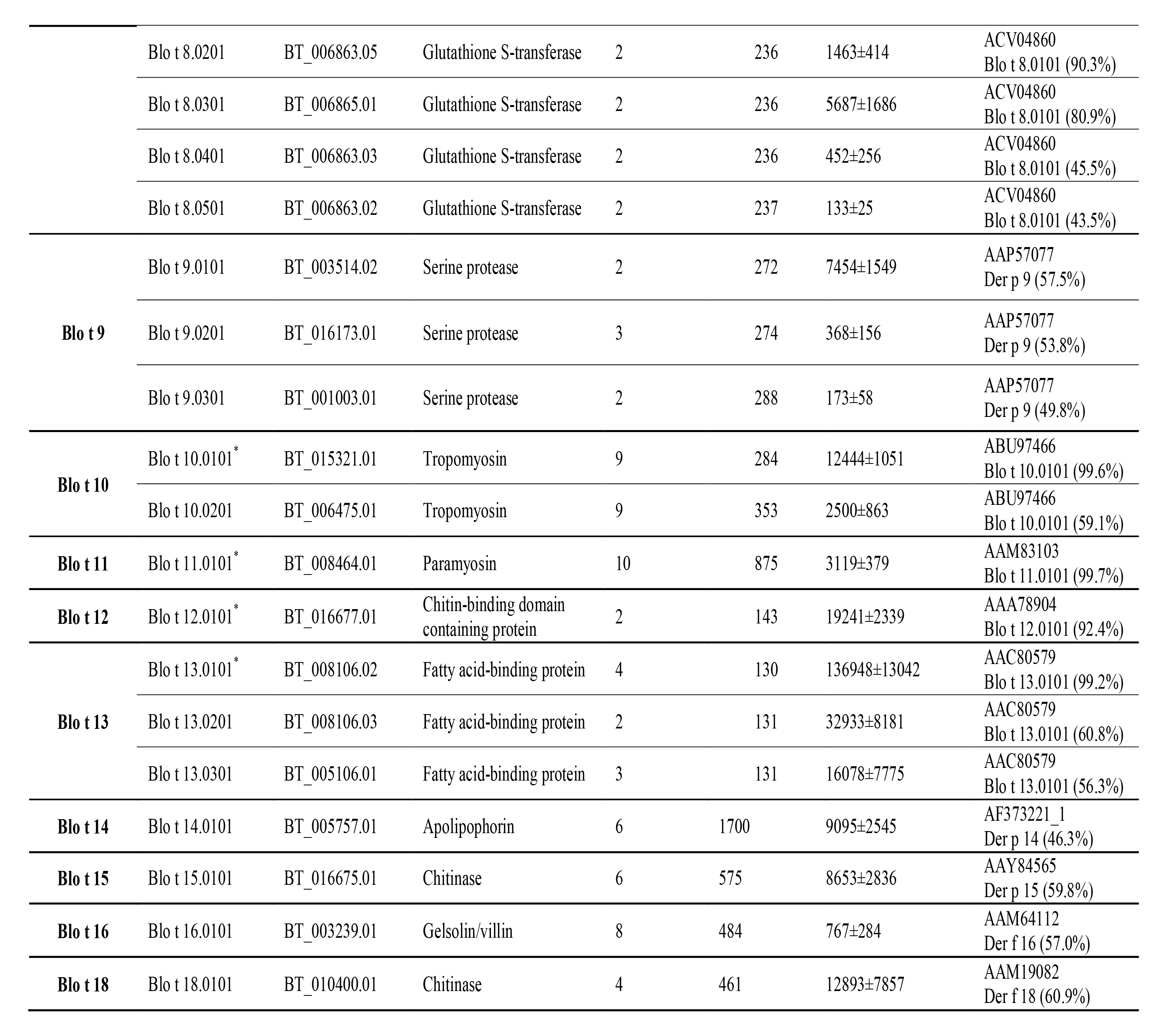
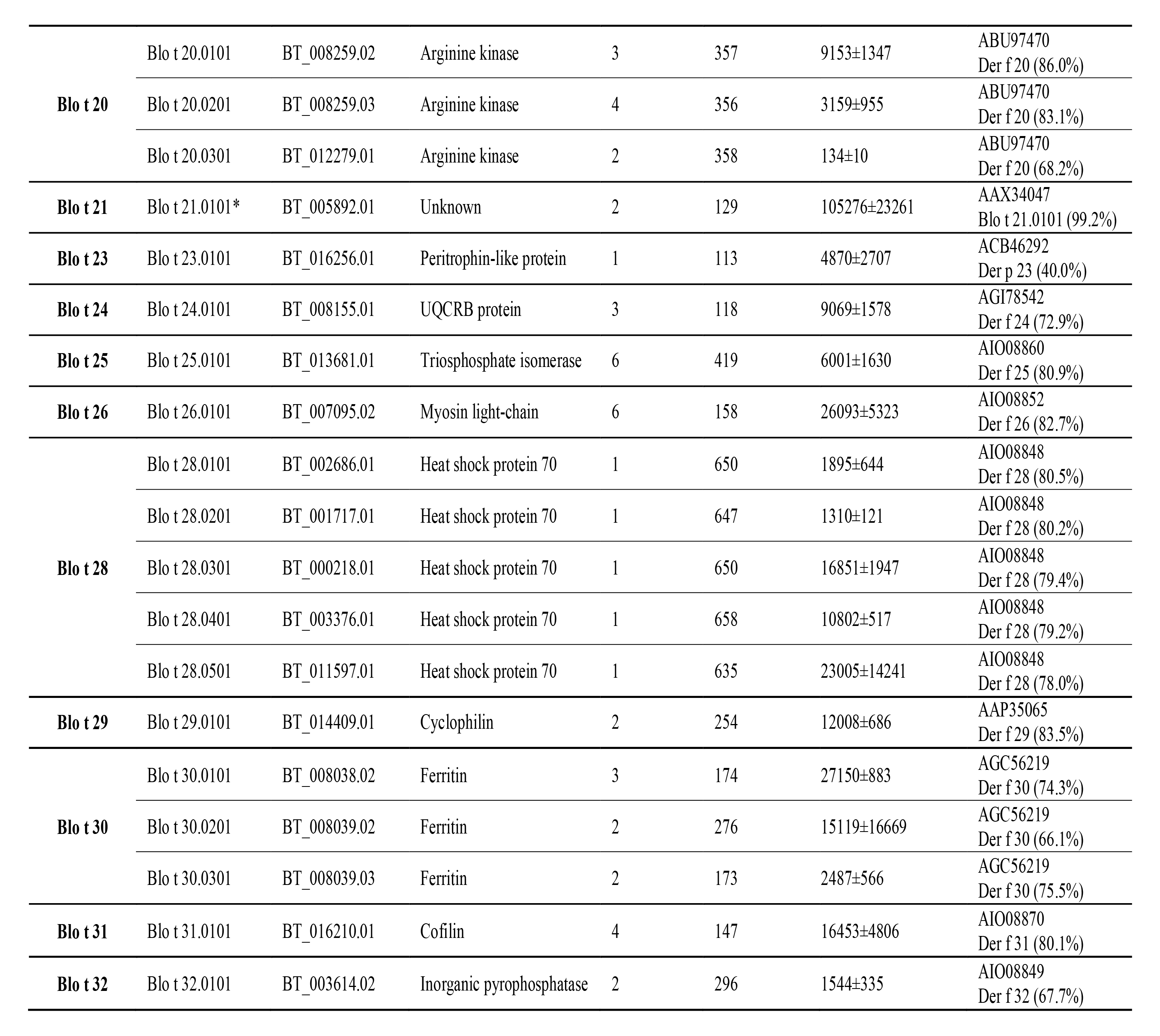
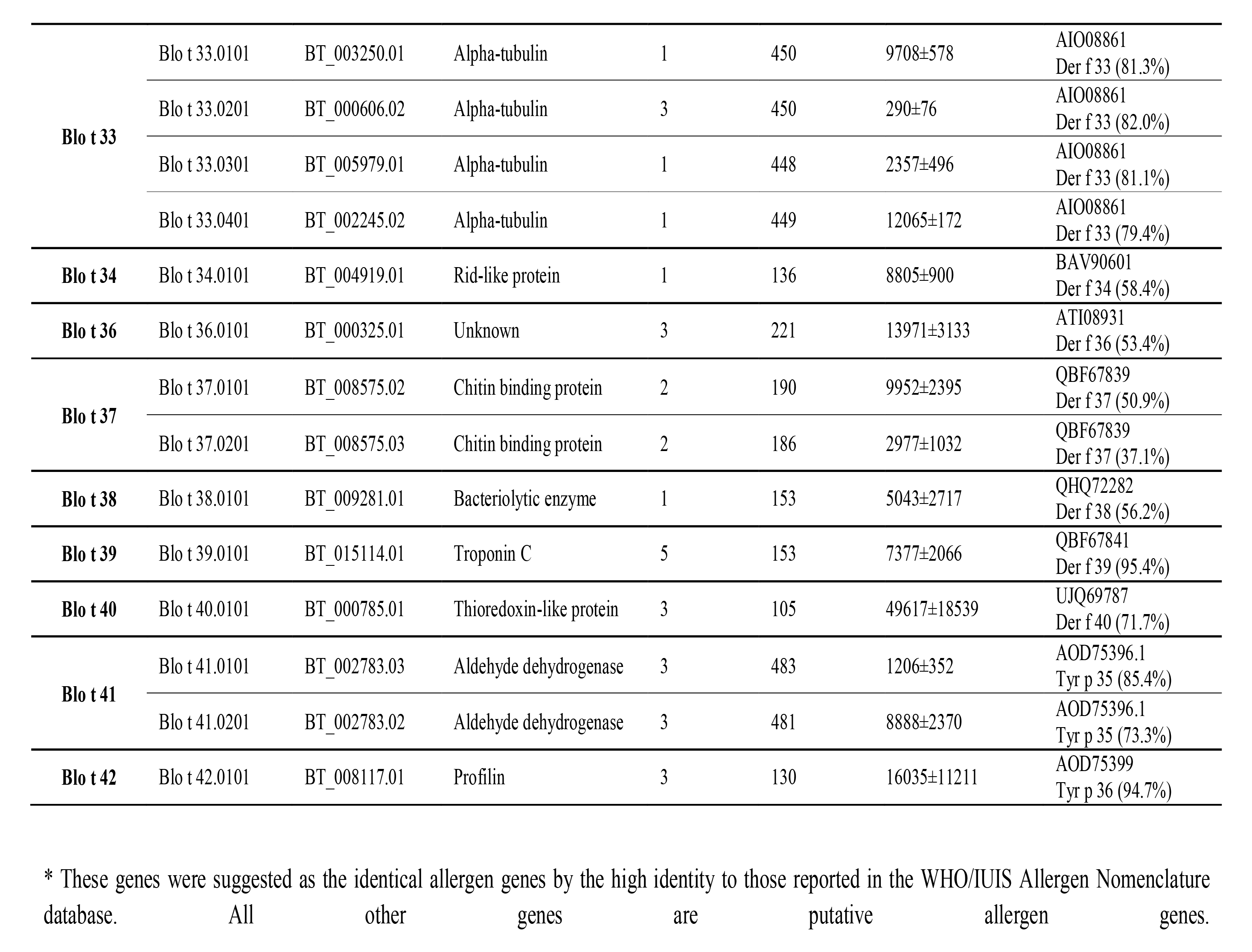
Summary of all allergen genes identified in the *B. tropicalis* genome.

### Quantification of gene expression level

To quantify the gene expression level of allergens, all the coding sequences (CDS) were collected and indexed using Salmon v0.12.0 (16) with the options ‘-i index --type quasi -k 31’. Then the transcriptome reads were mapped to the indexed CDS of all allergens using Salmon v0.12.0 (16) and the gene expression levels were represented by the transcript per million (TPM) values. The transcriptome data were obtained from pooled whole mite bodies and their NCBI SRA accessions SRR13742047, SRR13742048, SRR13742049 and SRR13742050 (Figure 1A). As for the gene expression of chitinases, the CDS of target genes were used as index. The bar chart and heatmaps were generated by the program GraphPad Prism v9.0.0.

**Figure 1.**
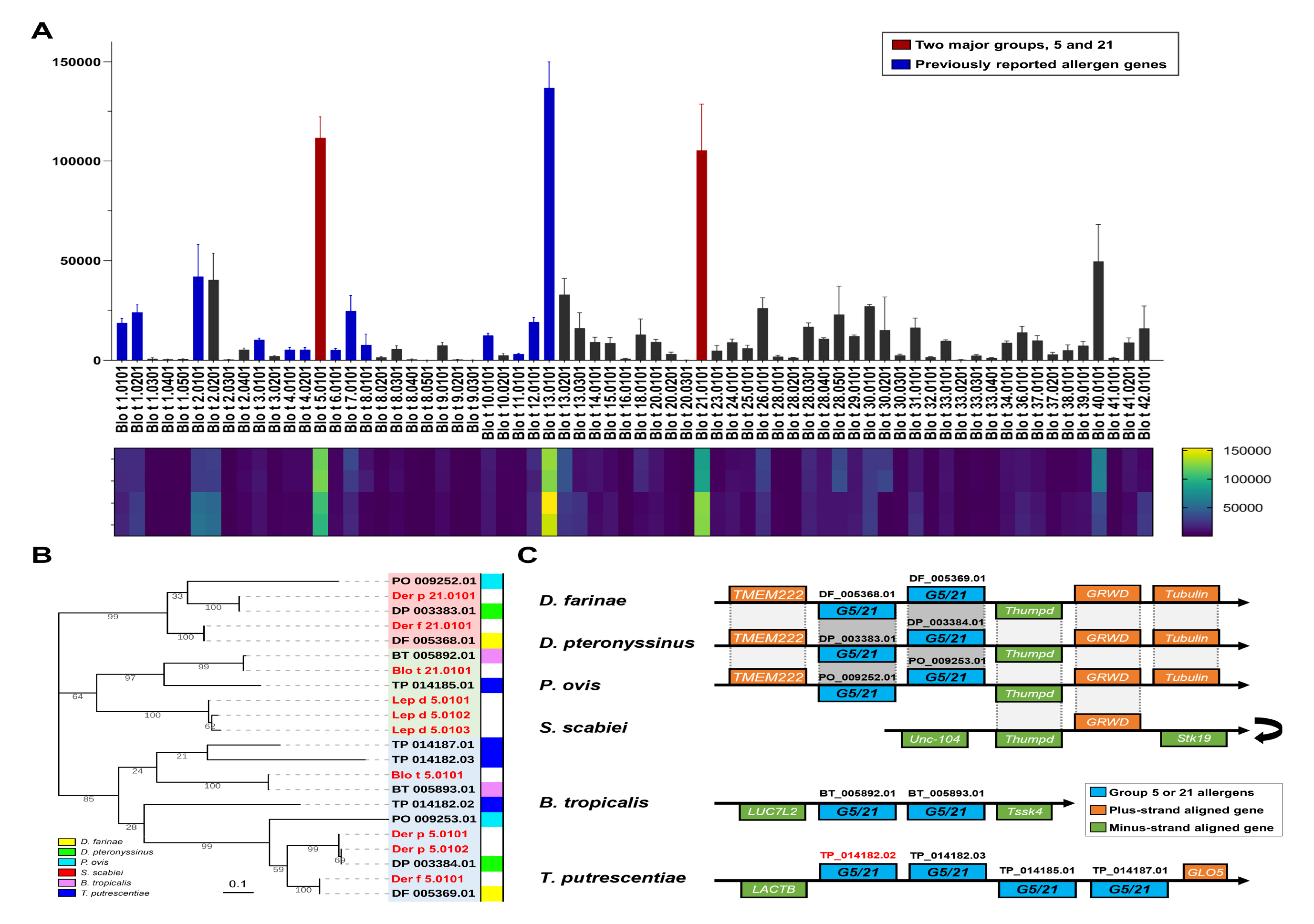
Genomic and transcriptomic analyses of *B. tropicalis* allergens. (A) Expression level of all the allergens genes identified in the *B. tropicalis* genome (Table 1). The gene expression level was represented by transcript per million (TPM) values. In the bar chart, error bar means standard deviation (SD). The coding sequences of all the allergen genes were used as index for read mapping of four transcriptome data, namely SRR13742048, SRR13742047, SRR13742050 and SRR13742049 (from top to bottom in the heatmap). (B) Phylogenetic analysis of the gene family of group 5 and 21 allergens in six astigmatic mites. All the allergens from the WHO/IUIS Allergen Nomenclature database were highlighted in red. (C) Gene synteny alignment of group 5 and 21 allergens in six astigmatic mites. A dimer gene of *T. putrescentiae* (TP_014182.02) was highlighted in red. G5/21, homologs of group 5 and 21 allergens. The turnover symbol means reverse complement.

### Cloning and expression of recombinant proteins

Protein and CDS sequences of specific allergens proteins of *B. tropicalis* were obtained after manual curation. The codon-optimized CDS sequences were synthesized and subcloned into plasmid vector (pET-28a+ for rBlo t 5, 18, 21, 23 and 26, pET-32a+ for rBlo t 12, 24, 25 and 31), and then transformed in *E. coli* cells, which was performed by Sangon Biotech (Shanghai, China). The *E. coli* cells were induced by isopropyl β-D-1-thiogalactopyranoside (final concentration: 1 mM) for protein expression and incubated at 16°C for 16 hours or 37°C for 4 hours. Proteins were extracted after cell ultrasonication, and 12% (w/v) SDS-PAGE analysis were performed to determine the protein solubility. The recombinant proteins were finally purified to over 90% by Ni^2+^ affinity chromatography using a fast protein liquid chromatography system. Four recombinant proteins, rBlo t 5, 12, 24 and 31, were soluble and purified from supernatant after centrifuge. The other five recombinant proteins were initially insoluble, then were denatured and purified from inclusion body pellets, and finally refolded in dialysis.

### Immunoassay of specific allergens

Allergen-specific IgE levels > 0.35 kUA/L were considered positive, which was further graded into classes 1 to 6 according to manufacturer’s instructions. For the ELISA experiments, serum samples were collected in Prince of Wales Hospital (Hong Kong) and obtained from 26 patients who presented with positive ImmunoCAP results (at least class 3) towards *B. tropicalis* (d201), and from 17 healthy individuals without mite sensitization as negative controls. The patients have at least one type of allergic disease in asthma, allergic rhinitis, and atopic dermatitis.

The allergenicity of the recombinant proteins was assessed by the enzyme-linked immunosorbent assay (ELISA). In the indirect sandwich ELISA, a 96-well microtiter plate was coated with 100 ul (2 ug/ml) of recombinant proteins per well in sodium bicarbonate coating buffer (0.1 mol/L, pH 9.5). The plate was washed with 0.05% (v/v) Tween-20 in PBS washing buffer for 4 times and then was blocked with 375 ul 3% (w/v) skim milk in PBST buffer for 30 minutes at 37□. Serum samples were diluted 1:4 in PBST containing 1% (w/v) BSA and 50 ul was added to each blocked well for 1 hr at room temperature. After washing, IgE antibodies were detected using biotinylated goat anti-human IgE (Vector, Burlingame, CA, USA) and streptavidin peroxidase (Sigma-Aldrich, St. Louis, MO, USA). The wells were then washed again with PBST buffer and incubated with 100 ul 3,3’,5,5’- Tetramethylbenzidine (Kirkegaard and Perry Laboratories, Gaithersburg, MD, USA) for 15 mins at room temperature. The reactions were stopped with the addition of 2M sulphuric acid (50 ul/well). The plates were then placed in the Benchmark Plus Microplate Spectrophotometer (BioRad, Hercules, CA, USA) and absorbance at 450 nm was recorded, where a high absorbance is indicative of a high concentration. The ELISA results were visualized and statistically analyzed in the program GraphPad Prism v9.0.0.

### Comparative analysis of gene families

All proteins of the six astigmatic mites, *D. farinae*, *D. pteronyssinus*, *Psoroptes* (*P.*) *ovis*, *Sarcoptes* (*S.*) *scabiei*, *B. tropicalis* and *Tyrophagus* (*T.*) *putrescentiae*, were searched by BLASTP v2.9.0 (12) with reference proteins at E-value cutoff of 1E-6. To collect all the genes in a family, the corresponding genes of astigmatic mite in Swiss-Prot database (17) (dated October 2021) or the WHO/IUIS allergen nomenclature database (4) (dated March 2022) were used for searching in the annotated protein sequences. After manual curation based on transcriptome data to confirm intron-exon split sites and filtering out annotated proteins with shorter than 50% average length of the gene family, all proteins of the six astigmatic mites identified in the target gene families were aligned and drawn into a phylogenetic tree. Sequence alignment was performed by CLUSTAL W (18) and MUSCLE (19) in MEGA v11.0.13 (20), and all phylogenetic trees were constructed by MEGA v11.0.13 (20) with maximum likelihood (ML) algorithm (21) in the Jones-Taylor-Thornton (JTT) model (22), 80% site coverage and 100 bootstrap replicates, and then edited by Interactive Tree of Life (iTOL) (23).

If two genes were located adjacently on genome and no other gene was located between them, they were considered as tandemly arrayed genes. If two genes were separated by no more than 10 genes, they were considered as proximally arrayed. Frequent tandemly arrayed genes were identified in gene families and connected with curve lines in phylogenetic trees edited by the online tool iTOL (23) with validated gene synteny information.

### Data availability

The genome and sequencing data of *B. tropicalis* are deposited in NCBI database under BioProject accession PRJNA702011. The *in silico* identified allergen sequences were uploaded in NCBI GenBank database under accessions OQ285922–OQ285989.

### Ethics approval

This study was approved by the institutional review board of Prince of Wales Hospital (CREC Ref. No.: 2018.663) for using the patient sera in ELISA experiments, and subjects and/or their parents have provided informed consent to participate.

## Results

### Genome-wide analysis of allergens

As an important HDM, *B. tropicalis* has only fourteen groups of allergens reported in the WHO/IUIS Allergen Nomenclature database (4). To fully understand the allergens of *B. tropicalis*, a genome-wide analysis was performed using the reported mite allergens as references to search in the high-quality genome (11). Thirty-seven allergen groups (up to Blo t 42) were identified and covered a comprehensive putative allergen profile of *B. tropicalis* (Table 1), including a wide range of homologs (or isoallergens). The reported group 19 allergen of *B. tropicalis*, Blo t 19 (GenBank accession: AHG97583), could not be found in the genome. In our naming rules, we used the first and last two digits after the decimal point to differentiate the different genes and isoforms, respectively. Multiple isoforms originating from the same gene locus were not included in this study, such as those caused by single-nucleotide polymorphism or alternative splicing. In our identification, group 2 and 12 allergens shared 93.8% and 92.4% identities with the reported allergens, respectively (Table 1). All other reported allergens in the database could be matched in our identified sequences with over 97% identity (Table 1). All these discrepancies could be explained by intraspecific differences.

The group 13 allergen, Blo t 13.0101, and the two major allergens of *B. tropicalis*, Blo t 5.0101 and 21.0101 (7, 24, 25), had significantly higher gene expression levels (Figure 1A). The high gene expression of the group 13 allergen was similarly found in the storage mite *T. putrescentiae* (26). The relatively high expression levels of the group 5 and 21 allergens may explain why they are major allergens of *B. tropicalis* but not the other mites. In the canonical HDMs *D. farinae* and *D. pteronyssinus*, we could not observe similar gene expression features (27).

Group 1 allergens of mites are characterized as cysteine proteases and are considered major allergens of HDMs but do not include *B. tropicalis*. Five homologs of group 1 allergens were identified in *B. tropicalis*, namely Blo t 1.0101, 1.0201, 1.0301, 1.0401 and 1.0501. Blo t 1.0101 and 1.0201 shared high similarities with the two reported Blo t 1 sequences (Table 1). All five homologs were proximally arrayed in the genome of *B. tropicalis* (Figure S1) and possibly originated from the duplication of one ancestral gene. Similarly, four homologs were observed in the group 2 allergen, in which Blo t 2.0201 and 2.0301 were tandemly arrayed (Figure S1). The best-matched group 2 allergen, Blo t 2.0101 (gene locus: BT_003416.02), shared only 93.8% identity with the reported Blo t 2.0101 and the difference was mainly caused by a mismatch region on the N-terminus (Figure S2). Along with groups 22 and 35 allergens of mites (including Der f 22, Der f 35 and Der p 22(28)), group 2 allergens belong to the Niemann-Pick protein type C2 (NPC2) family and have been well analyzed (26).

Group 3, 6 and 9 allergens are all members of the serine protease family (5), in which massive tandem gene duplications of *B. tropicalis* occurred (11). According to the adapted phylogenetic tree (11), the three serine protease allergens were located in distinct clusters (Figure S3). Group 3 allergens were close to the species-specific inactive serine proteases of *B. tropicalis* and *S. scabiei*, while group 6 and 9 allergens had no close homolog from the two parasitic mites *P. ovis* and *S. scabiei* (Figure S3).

At least two highly identical genes encoding alpha amylase, were identified as single-exon genes, and shared over 99% identity with the reported group 4 allergen, Blo t 4 (GenBank accession: AAQ24543). Because these two homologs of Blo t 4 are located on different scaffolds in different gene syntenies, we determined that they were generated via a recent transposable duplication. Initially, four homologous genes of Blo t 4 were identified in *B. tropicalis* (11). However, two low-quality genes were discarded because one gene was located on a short scaffold encoding only one gene, while the other gene on a short scaffold shared highly conserved synteny with BT_008084.02 and was considered a heterozygous gene.

For the major allergen groups 5 and 21 of *B. tropicalis* (7, 24, 25), we noticed that Blo t 5.0101 and 21.0101 are tandemly arrayed homologs that share 41.9% identity. To further explore the divergence of this gene family (G5/21), we collected all the genes of six mites and performed phylogenetic analysis (Figure 1B). Two major allergens, groups 5 and 21 of *B. tropicalis*, were compared with those of the other five astigmatic mites (11) (Figure 1B). In *D. farinae*, *D. pteronyssinus* and *P. ovis*, the two tandemly arrayed G5/21 genes were located on opposite strands and shared identical gene synteny (Figure 1C), which supports the close phylogenetic relationship among the three species (11). For the skin-burrowing mite *S. scabiei*, the absence of the G5/21 gene may be caused by complete gene decay. In two canonical storage mites, *B. tropicalis* and *T. putrescentiae*, the gene syntenies surrounding G5/21 genes are totally different from those in other species (Figure 1C). Unlike *D. farinae*, *D. pteronyssinus* and *P. ovis*, the two G5/21 genes of *B. tropicalis* are located on the same strand. For *T. putrescentiae*, four tandemly arrayed G5/21 genes, including a dimer gene TP_014182.02, were identified (26). Both group 5 and 21 allergens were highly expressed in *B. tropicalis* (Figure 1A) but expressed at low levels in *T. putrescentiae* (26). We suggest that group 5 and 21 allergens of mites are homologous genes that originated from an ancestral gene.

Only one homolog of the group 7 allergen was identified, while up to five tandemly arrayed homologs of group 8 allergens were found in *B. tropicalis* (Table 1, Figure S1). Group 8 allergens of mites were characterized as glutathione-S-transferase (5), of which the gene family variation has been well explored (26). Two tropomyosins and one paramyosin were identified in group 10 and 11 allergens, respectively (Table 1). Notably, the group 12 allergen, a protein containing a chitin-binding domain that has not been reported in other HDMs, could be identified in *B. tropicalis* (gene locus: BT_016677.01) but presented only 92.4% identity to that reported for Blo t 12.0101 (GenBank accession: AAA78904, Table 1). In group 13 allergens, three homologs were found, including two tandemly arrayed genes (Table 1, Figure S1).

Moreover, a wide range of newly identified allergen groups were identified in *B. tropicalis* (Table 1). All these newly predicted allergens of *B. tropicalis* exhibit high similarities (>45% identity) to those of *D. pteronyssinus* (prefix: Der p) or *D. farinae* (prefix: Der f), and no homolog of the group 27 allergen was identified. Along with group 2 allergens, group 22 and 35 allergens are homologs in the NPC2 family (26) and were not identified in *B. tropicalis*.

### Comparative analysis of chitin-binding allergens

Chitinases and other chitin-binding domain containing proteins have been reported to play important roles in human allergic disease (29, 30). Groups 12, 23 and 37 allergens of mites contain chitin-binding domains, while groups 15 and 18 allergens are characterized as chitinases (5).

Group 12 allergen was only reported in *B. tropicalis* (31) in the WHO/IUIS allergen nomenclature database. The reported Blo t 12 (GenBank accession: AAA78904) was from mites collected in Colombia, while the *B. tropicalis* mite in this study originated from Singapore (NCBI BioSample: SAMN17935020). Therefore, it is reasonable that our identified Blo t 12 is nearly identical (only one different amino acid) to that from mites collected in Singapore (32) but shares only 92.4% identity with the reported Blo t 12. In sequence alignment, we observed that the major difference was the consequence of a regional frame-shift mutation (block 2, Figure S4). To explore the genomic specificity of the group 12 allergen, gene synteny analysis was performed among six astigmatic mites (Figure 2A). The group 12 allergen gene of *B. tropicalis*, Blo t 12, was located adjacently with a chitinase gene that shared as high as 59.8% identity in protein sequence with the group 15 allergen of *D. pteronyssinus* and identified as Blo t 15 (Table 1). Therefore, Blo t 12 and 15 were tandemly arrayed and shared high conservation on their C-terminal sequences (chitin-binding domain). (Figure S5). Group 23 was reported in *D. farinae* and *D. pteronyssinus* (33, 34). We identified a homolog of group 23 allergen in the annotated genome of *B. tropicalis*, Blo t 23, that shares 40.0% identity with Der p 23 (GenBank accession: ACB46292). Despite the low similarity, the chitin-binding domains of Blo t 23 and Der p 23 were highly conserved (Figure S6).

**Figure 2.**
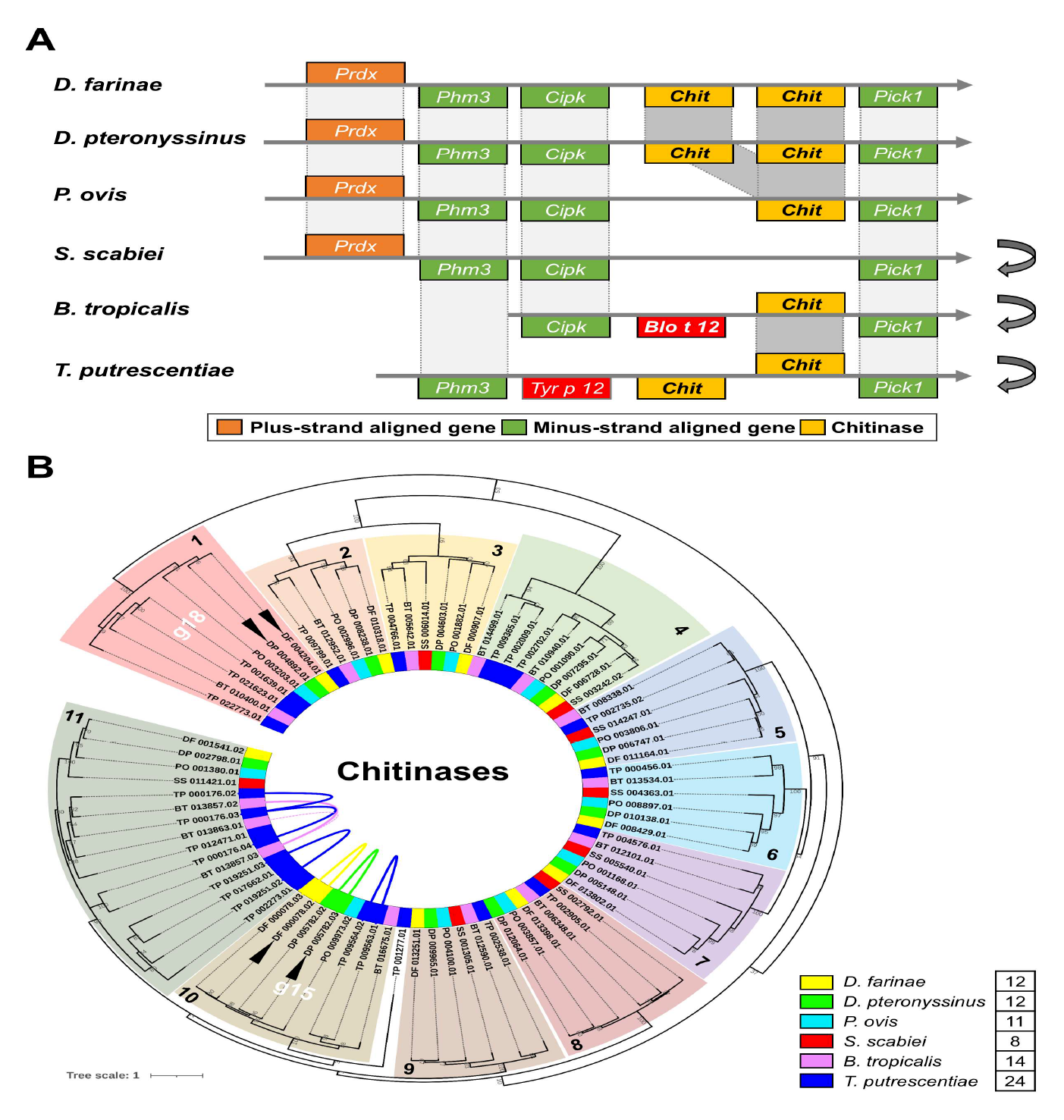
Comparative analysis of chitin-binding allergens. (A) Gene synteny alignment of the group 12 allergen loci in six astigmatic mites. The group 12 allergen homologs of *B. tropicalis* and *T. putrescentiae* and the chitinase genes were highlighted in red and yellow boxes, respectively. The turnover symbol means reverse complement. None of group 12 allergen homolog or chitinase was identified in the locus of *S. scabiei*. (B) Phylogenetic analysis of chitinase genes in six astigmatic mites. Eleven clusters were classified, including the Cluster 1 and 10 identified as the group 18 and 15 allergens, respectively. Four reported allergens in the WHO/IUIS Allergen Nomenclature database were marked by black triangles. Tandemly arrayed genes and proximally arrayed genes (separated by no more than 10 genes) were connected by curved solid lines and dotted lines, respectively.

Group 15 and 18 allergens of mites are both identified as chitinases. All the chitinase genes of the six astigmatic mites were divided into eleven clusters in the phylogenetic tree (Figure 2B). Tandem gene duplications were observed in Clusters 10 and 11(Figure 2B). Clusters 2 and 3 were horizontally transferred genes, as previously reported (11). The chitinases in the two distinct Clusters 1 and 10 were identified as group 18 and 15 allergens of mites, respectively (Figure 2B). Interestingly, both group 15 and 18 allergens decayed in the genome of *S. scabiei* (Figure 2B). The gene synteny alignment supported the gene decay of the group 15 allergen in *S. scabiei* (Figure 2A). Both group 15 and 18 allergens are highly conserved among astigmatic mites in protein sequences (Figures S7 and S8). In gene expression analysis, we observed the consistently high expression of group 15 and 18 allergens in astigmatic mites (Figure S9). In particular, the chitinases in Cluster 11 were highly expressed in two parasitic mites, *P. ovis* and *S. scabiei* (Figure S9).

### Immunoassay of recombinant allergen proteins

Although our genome-wide analysis revealed a comprehensive allergen profile of *B. tropicalis* (Table 1), serological evidence is still needed to confirm the allergenicity of many novel allergens. To verify and evaluate the allergenicity of the proposed novel allergens, their recombinant proteins were expressed and subjected to ELISA experiment using patient serum samples. Serum samples were collected in Hong Kong from allergic patients for testing and healthy individuals as controls (Table S1). When at least class 3 allergy (over 3.5 kUA/L of IgE) was considered positive, 58% (29/50) of those tested positive for HDM (Der p 1, *D. pteronyssinus*) were identified to be positive for *B. tropicalis*, while only one sample (BC0699 out of 20 samples) was negative for HDM (class 0) but positive for *B. tropicalis* (class 4). The sensitization rate of those serum samples was apparently lower than that in Singapore where the prevalence of *B. tropicalis* could be up to 97.5% in allergic children (35).

In subsequent ELISA experiments, 26 serum samples were collected from allergic patients who tested positive for *B. tropicalis* (BT-positive). Nine recombinant proteins of *B. tropicalis* were expressed and purified (Figure S10), including three reported allergens (rBlo t 5, 21 and 12) and six novel proposed allergens (rBlo t 18, 23, 24, 25, 26 and 31). The absorbance values at 450 nm of recombinant proteins represented the IgE levels of specific allergens (Table S2, Figure 3 and S11). The major allergens rBlo t 5 and 21 were used as positive controls (Figure 3A and B). According to the allergen-specific IgE levels in BT-allergic sera, rBlo t 18, 23 and 26 were verified as novel allergens of *B. tropicalis* (Figure 3C-E), while rBlo t 24, 25 and 31 were not confirmed to be novel allergens, because of their low IgE levels (Figure S11). Unexpectedly, rBlo t 12 presented significantly higher IgE levels in BT-allergic sera, but the low absorbance values (all below 0.15) could not well confirm its allergenicity (Figure S11A), which was similarly observed in rBlo t 24 (Figure S11B). The IgE levels of two major allergens, rBlo t 5 and 21 (Figure 3A and B), were highly dispersed with high maximum values (above 1.0) and standard deviations (SDs), which affected the significance levels (*P* values). Similar features were observed in three novel allergens, rBlo t 18, 23 and 26 (Figure 3C-E). Intriguingly, regression analysis revealed good correlations among the IgE levels of rBlo t 5, 21, 23 and 26, but not rBlo t 18, and suggested the cosensitization of these allergens (Figure S12).

**Figure 3.**
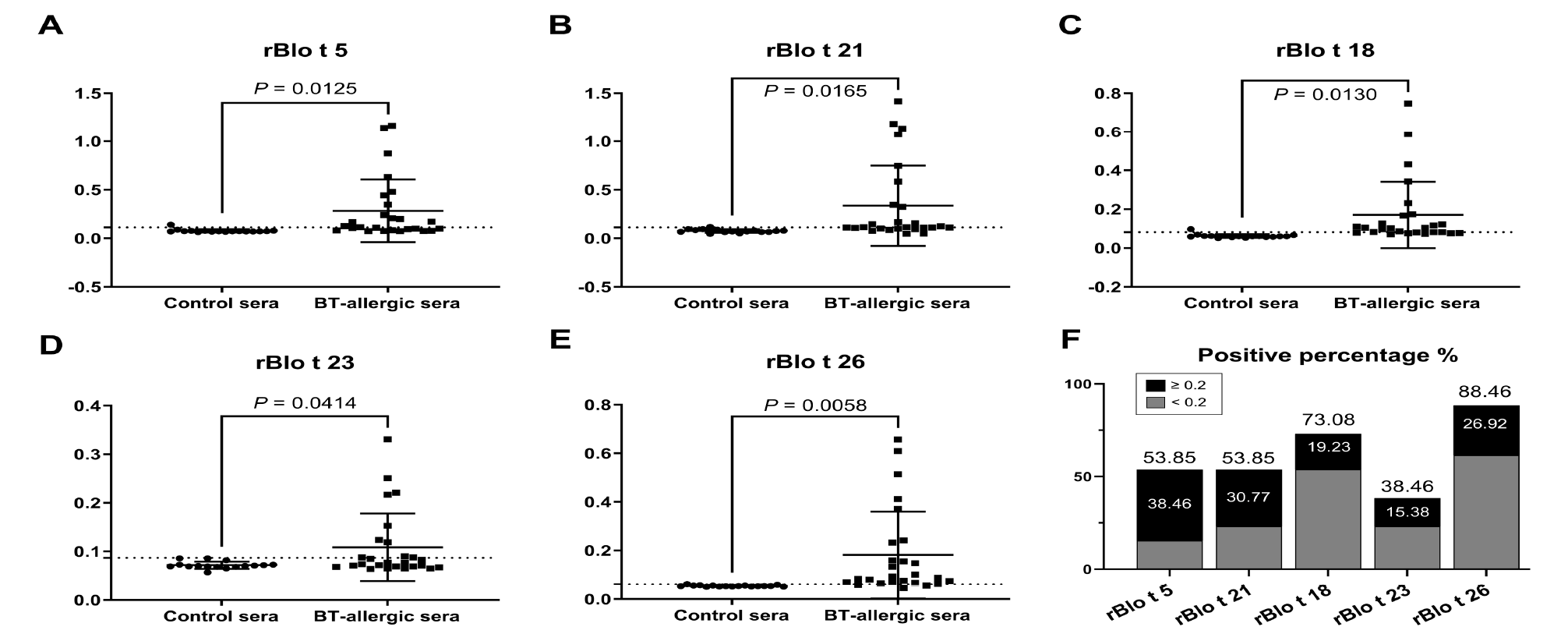
ELISA results of recombinant allergen proteins of *B. tropicalis*. The IgE antibody binding to recombinant proteins of *B. tropicalis* was evaluated by ELISA using serum samples from 26 patients positive for *B. tropicalis* (BT-allergic sera) for testing and 17 healthy individuals as control sera (Table S1). The absorbance values at 450 nm of recombinant proteins were recorded to represent the IgE levels of specific allergens. The lines in dot plots were set as the mean with a standard deviation (SD) and the dotted lines indicated the cutoff value, mean + 2*SD of the healthy controls. The allergenicity was evaluated for five recombinant proteins as follows: (A) rBlo t 5, (B) rBlo t 21, (C) rBlo t 18, (D) rBlo t 23, (E) rBlo t 26. Unpaired two-tailed t-test P-values indicate statistical significance (all below 0.05). (F) Positive percentage of five recombinant proteins for the allergens of *B. tropicalis*. The absorbance values equal to or higher than 0.2 were regarded as high IgE and allergy levels and their percentages were highlighted in columns.

For the sensitization rates of specific allergens (Figure 3F), unexpectedly, rBlo t 26 and 18 presented high positive percentages of 88.46% and 73.08%, respectively, while two major allergens rBlo t 5 and 21 demonstrated only 53.85%, and the chitin-binding allergen rBlo t 23 showed low sensitization rates 38.46%. The prevalence observed in this batch of Hong Kong sera was totally different from that found in Singapore samples, which showed the highest positivity to the two major allergens Blo t 5 and 21 (7). When we regarded absorbance values equal to or higher than 0.2 as high IgE and allergy levels, two major allergens, rBlo t 5 and 21, demonstrated higher percentages of 38.46% and 30.77% (Figure 3F).

## Discussion

With the advent of omics era, a series of genome-based approaches have been applied to the study of allergens and mite allergies (11, 27, 36–41). Based on the high-quality genome of the dust mite *B. tropicalis*, our genome-wide analysis identified a comprehensive putative allergen profile of thirty-seven allergen groups (up to Blo t 42), which covered a wide range of allergen homologs. The specific allergen Blo t 12 has been well confirmed, while Blo t 19 was considered nonexistent in the genome. Considering group 19 allergen could not been identified in *D. farinae*, we speculated that the allergen Blo t 19 was falsely reported (42). Two cysteine proteases of *B. tropicalis* have been identified as isoallergens in group 1. In the allergen profile, many more homologs of *B. tropicalis* allergens were identified and proposed to be allergens (Table 1), including up to five homologs of group 1 allergens. Consistent with a previous report (11), tandem gene duplication could cause gene family expansion and result in multiple homologs (Figure S1), such as five tandemly arrayed homologous genes of Blo t 8 (glutathione S-transferase). Transcriptomic analysis revealed interesting gene expression levels of the *B. tropicalis* allergens. Unexpectedly, the Blo t 13 homolog, Blo t 13.0101, presented the highest expression level (Figure 1A), which was similarly observed in the storage mite *T. putrescentiae* (26). After Blo t 13.0101, two major allergens of *B. tropicalis*, Blo t 5 and 21, were the two most highly expressed allergens (Figure 1A), which may be related to why they were the major allergens. Blo t 13 was not identified to be major allergen (43), possibly because it could only be found in mite bodies rather than fecal pellets (44). The transcriptome data and expression levels are highly variable with geographical location, mite collection and culture conditions, especially when Blo t 2 was identified to be the most abundant allergen in a previous report (45). Therefore, multiple sampling is necessary in future allergen abundance analysis of mites.

To better understand the complicated allergen profile, more comparative analysis was performed for important allergen groups. A wide range of tandem duplications were found in the expansion and evolution of many allergen groups. In particular, the two major allergens of *B. tropicalis*, Blo t 5 and 21, were identified to be tandemly arrayed homologs. In the gene family of group 5 and 21 allergens, two tandemly arrayed homologs were identified in other astigmatic mites, including *D. farinae*, *D. pteronyssinus* and *P. ovis*. Four homologs were found in the storage mite *T. putrescentiae*, while none was observed in the skin-borrowing mite *S. scabiei* (Figure 1C). Not only the specific Blo t 12 but also group 15, 18 and 23 allergens of mites contain chitin-binding domains. Gene synteny and sequence alignment suggested that the group 12 allergens were specific to canonical storage mites (including *B. tropicalis* and *T. putrescentiae*) and derived from the group 15 allergen (chitinase). Comparative analysis of chitinases further revealed the divergence of group 15 and 18 allergens that are especially allergenic in dogs. Despite the absence of aligned gene synteny, Blo t 23 was identified with a conserved chitin-binding domain shared with Der p 23. Such a wide range of chitin-binding allergens would be an interesting topic of mite allergy and deserves further exploration.

In the ELISA test of recombinant proteins (Figure 3), allergenicity was assessed for five specific allergens of *B. tropicalis*, including three novel allergens, Blo t 18, 23 and 26. In particular, Blo t 18 and 26 presented not only high positivity but also a relatively high proportion of high allergy levels, which suggests that they are important novel allergens of *B. tropicalis*. The high proportion of high allergy levels to rBlo t 5 and 21 indicated the importance of these two major allergens. The sensitization rates of specific allergens in this batch of Hong Kong sera demonstrated totally distinct features from those observed in Singapore samples with obviously high sensitization rates of Blo t 5 and 21 (7), which indicated the geographical difference in *B. tropicalis* allergy. Considering that nearly all the samples (BT-positive) were also positive for *D. pteronyssinus* (Table S1), we cannot exclude the influence of cross-reactivity in the ELISA results. A major limitation of this study is that only nine recombinant proteins were expressed.

In summary, this genome-wide analysis shed light on the allergen profile in *B. tropicalis*, the comparative analysis revealed insights into the divergence of allergen gene families, and the immunoassay experiments of recombinant proteins provided serological evidence for novel allergens. Taken together, this comprehensive analysis of *B. tropicalis* paves the way to the CRD of mite allergy.

## Abbreviations

HDM: house dust mite;
ELISA: enzyme-linked immunosorbent assay;
NPC2: Niemann-Pick protein type C2;
CRD: component-resolved diagnosis

## Acknowledgments

We would like to thank the staff members in the Siriraj Dust Mite Center for Services and Research, Siriraj Hospital, Bangkok, Thailand, for the mite culturing work.

This work was made possible by grants:

1. General Research Fund from Research Grants Council of Hong Kong (Reference numbers: 464710, 475113, 14119219, 14119420, 14175617)
2. Health and Medical Research Fund from Food and Health Bureau of Hong Kong (Reference numbers: 06171016, 07181266)
3. Continuation project of Joint Research Fund for Overseas Chinese Scholars and Scholars in Hong Kong and Macao Young Scholars (Reference number: 31729002)
4. National Natural Science Foundation of China (Reference numbers: 81971514, 82073950)
5. Shenzhen Science and Technology Plan Project (Reference number: GJHZ20190822095605512, SGDX20201103095609027)

## Supplementary data

Tables S1–2

Figures S1–12

