## Supplementary Materials for "Genomic Analysis of *Blomia tropicalis* Identifies Novel Allergens for Component-Resolved Diagnosis of Mite Allergy"

**Table S1. Basic information of the serum samples used in ELISA test**

Serum samples were collected from 26 mite-sensitized allergic patients and 17 healthy individuals without mite sensitization (mean age, 12.0 years old; range, 5.8-18.6). The patients have at least one type of allergic disease in asthma, allergic rhinitis and atopic dermatitis. Allergy levels to house dust mite (Der p 1, *D. pteronyssinus*) were measured by fluorescent enzyme immunoassay AutoCAP (Phadia AB, Uppsala, Sweden), and those to *Blomia tropicalis* were evaluated by ImmunoCAP (Thermo Fisher Scientific, USA) using the allergen d201. Data with specific IgE concentration > 0.35 kUA/L were considered positive, which was further graded into classes 1 to 6 according to manufacturer's instructions.

| Sample ID | Age (y/o) | Allergy level to house dust mite (class) | Allergy level to *Blomia tropicalis* | |
|---|---|---|---|---|
| | | | Concentration (kUA/l) | Class |
| **Allergic patients** | | | | |
| A753 | 9.9 | 5 | 4.22 | 3 |
| A756 | 17.8 | 5 | 8.96 | 3 |
| A760 | 12 | 5 | 9.73 | 3 |
| BC0050 | 9.4 | 6 | 9.56 | 3 |
| BC0071 | 13.3 | 5 | 11.4 | 3 |
| BC0111 | 8.9 | 4 | 11 | 3 |
| BC0118 | 10.4 | 5 | 26.3 | 4 |
| BC0133 | 13.5 | 6 | 6.95 | 3 |
| BC0146 | 13.1 | 4 | 9.63 | 3 |
| BC0151 | 8.2 | 6 | 38.8 | 4 |
| BC0154 | 5.8 | 6 | 6.26 | 3 |
| BC0157 | 13.5 | 6 | 4.47 | 3 |
| BC0160 | 13 | 5 | 13.8 | 3 |
| BC0161 | 14.2 | 6 | 6.61 | 3 |
| BC0162 | 14 | 5 | 3.71 | 3 |
| BC0171 | 9.4 | 6 | 29.4 | 4 |
| BC0172 | 9.1 | 6 | 66.9 | 5 |
| BC0177 | 13 | 5 | 9.1 | 3 |
| BC0182 | 10.5 | 6 | 14.9 | 3 |
| BC0183 | 14.3 | 4 | 31.3 | 4 |
| BC0191 | 18.6 | 6 | 12.7 | 3 |
| BC0194 | 11 | 6 | >100 | 6 |
| BC0465 | 9.7 | 6 | 3.66 | 3 |
| BC0482 | 17 | 6 | 52.7 | 5 |
| BC0507 | 13.9 | 5 | 32.9 | 4 |
| BC0699 | 8.1 | 0 | 26.6 | 4 |
| **Healthy control** | | | | |
| BC0062 | 7.2 | 0 | 0.02 | 0 |
| BC0080 | 6.9 | 0 | 0.05 | 0 |
| BC0127 | 13.2 | 0 | 0.01 | 0 |
| BC0137 | 13.0 | 0 | 0.01 | 0 |
| BC0149 | 8.7 | 0 | 0.03 | 0 |
| BC0178 | 11.4 | 0 | 0.01 | 0 |
| BC0215 | 14.2 | 0 | 0.01 | 0 |
| BC0217 | 14.5 | 0 | 0.02 | 0 |
| BC0228 | 14.5 | 0 | 0.01 | 0 |
| BC0231 | 14.8 | 0 | 0.16 | 0 |
| BC0235 | 14.1 | 0 | 0.01 | 0 |
| BC0237 | 14.9 | 0 | 0.03 | 0 |
| A747 | 7.2 | 0 | 0.02 | 0 |
| A754 | 16.5 | 0 | 0.02 | 0 |
| A757 | 8.8 | 0 | 0.01 | 0 |
| A766 | 15.7 | 0 | 0.01 | 0 |
| A767 | 7.7 | 0 | 0.01 | 0 |

**Table S2. Allergen-specific IgE levels represented by absorbance value at 450 nm**

| Sample ID | rBlo t 5 | rBlo t 21 | rBlo t 12 | rBlo t 18 | rBlo t 23 | rBlo t 24 | rBlo t 25 | rBlo t 26 | rBlo t 31 |
|---|---|---|---|---|---|---|---|---|---|
| **Allergic patients** | | | | | | | | | |
| A753 | 0.171 | 0.155 | 0.118 | 0.174 | 0.065 | 0.073 | 0.058 | 0.159 | 0.042 |
| A756 | 0.095 | 0.099 | 0.087 | 0.08 | 0.071 | 0.083 | 0.058 | 0.148 | 0.053 |
| A760 | 0.106 | 0.114 | 0.127 | 0.231 | 0.069 | 0.095 | 0.053 | 0.088 | 0.055 |
| BC0050 | 0.076 | 0.346 | 0.092 | 0.077 | 0.077 | 0.085 | 0.058 | 0.155 | 0.069 |
| BC0071 | 0.109 | 0.746 | 0.125 | 0.587 | 0.088 | 0.075 | 0.056 | 0.371 | 0.048 |
| BC0111 | 0.443 | 0.325 | 0.107 | 0.167 | 0.119 | 0.09 | 0.059 | 0.242 | 0.059 |
| BC0118 | 1.165 | 1.135 | 0.094 | 0.07 | 0.217 | 0.094 | 0.057 | 0.412 | 0.064 |
| BC0133 | 0.2 | 0.166 | 0.112 | 0.104 | 0.088 | 0.112 | 0.057 | 0.133 | 0.058 |
| BC0146 | 0.077 | 0.078 | 0.084 | 0.076 | 0.068 | 0.068 | 0.052 | 0.074 | 0.055 |
| BC0151 | 1.142 | 1.073 | 0.098 | 0.079 | 0.251 | 0.08 | 0.063 | 0.514 | 0.079 |
| BC0154 | 0.1 | 0.146 | 0.112 | 0.431 | 0.071 | 0.104 | 0.056 | 0.075 | 0.049 |
| BC0157 | 0.1 | 0.111 | 0.114 | 0.747 | 0.067 | 0.075 | 0.055 | 0.045 | 0.044 |
| BC0160 | 0.208 | 0.124 | 0.113 | 0.097 | 0.065 | 0.626 | 0.055 | 0.058 | 1.091 |
| BC0161 | 0.24 | 0.109 | 0.102 | 0.343 | 0.064 | 0.065 | 0.053 | 0.083 | 0.077 |
| BC0162 | 0.347 | 0.049 | 0.098 | 0.116 | 0.085 | 0.084 | 0.056 | 0.1 | 0.053 |
| BC0171 | 0.085 | 0.114 | 0.068 | 0.105 | 0.072 | 0.123 | 0.057 | 0.093 | 0.061 |
| BC0172 | 0.074 | 0.111 | 0.108 | 0.076 | 0.083 | 0.097 | 0.064 | 0.068 | 0.082 |
| BC0177 | 0.115 | 0.101 | 0.075 | 0.073 | 0.074 | 0.086 | 0.052 | 0.08 | 0.064 |
| BC0182 | 0.082 | 0.095 | 0.105 | 0.083 | 0.124 | 0.067 | 0.065 | 0.065 | 0.069 |
| BC0183 | 0.631 | 0.584 | 0.12 | 0.127 | 0.153 | 0.088 | 0.064 | 0.232 | 0.071 |
| BC0191 | 0.126 | 0.111 | 0.107 | 0.122 | 0.069 | 0.074 | 0.059 | 0.063 | 0.087 |
| BC0194 | 0.074 | 0.107 | 0.097 | 0.082 | 0.065 | 0.102 | 0.054 | 0.056 | 0.055 |
| BC0465 | 0.076 | 0.048 | 0.093 | 0.11 | 0.077 | 0.089 | 0.059 | 0.069 | 0.042 |
| BC0482 | 0.874 | 1.418 | 0.096 | 0.086 | 0.331 | 0.085 | 0.067 | 0.61 | 0.056 |
| BC0507 | 0.479 | 1.184 | 0.104 | 0.102 | 0.221 | 0.095 | 0.056 | 0.657 | 0.083 |
| BC0699 | 0.165 | 0.088 | 0.1 | 0.082 | 0.09 | 0.088 | 0.057 | 0.067 | 0.1 |
| **Healthy control** | | | | | | | | | |
| BC0062 | 0.07 | 0.065 | 0.066 | 0.053 | 0.069 | 0.067 | 0.052 | 0.052 | 0.102 |
| BC0080 | 0.09 | 0.066 | 0.06 | 0.06 | 0.086 | 0.075 | 0.053 | 0.056 | 0.081 |
| BC0127 | 0.074 | 0.071 | 0.071 | 0.056 | 0.068 | 0.065 | 0.054 | 0.053 | 0.068 |
| BC0137 | 0.067 | 0.076 | 0.076 | 0.057 | 0.07 | 0.067 | 0.053 | 0.054 | 0.062 |
| BC0149 | 0.077 | 0.066 | 0.074 | 0.06 | 0.073 | 0.063 | 0.055 | 0.053 | 0.059 |
| BC0178 | 0.071 | 0.082 | 0.079 | 0.059 | 0.072 | 0.074 | 0.078 | 0.052 | 0.057 |
| BC0215 | 0.078 | 0.05 | 0.068 | 0.058 | 0.069 | 0.054 | 0.049 | 0.05 | 0.053 |
| BC0217 | 0.073 | 0.094 | 0.05 | 0.062 | 0.065 | 0.053 | 0.052 | 0.053 | 0.056 |
| BC0228 | 0.069 | 0.07 | 0.062 | 0.06 | 0.073 | 0.063 | 0.053 | 0.062 | 0.063 |
| BC0231 | 0.064 | 0.08 | 0.058 | 0.097 | 0.068 | 0.066 | 0.049 | 0.051 | 0.055 |
| BC0235 | 0.07 | N. A.* | 0.075 | 0.061 | 0.086 | 0.072 | 0.088 | 0.059 | 0.074 |
| BC0237 | 0.074 | 0.118 | 0.081 | 0.06 | 0.071 | 0.067 | 0.058 | 0.056 | 0.083 |
| A747 | 0.072 | 0.086 | 0.071 | 0.067 | 0.069 | 0.066 | 0.054 | 0.057 | 0.066 |
| A754 | 0.073 | 0.095 | 0.087 | 0.062 | 0.07 | 0.063 | 0.054 | 0.054 | 0.062 |
| A757 | 0.075 | 0.067 | 0.114 | 0.069 | 0.082 | 0.066 | 0.053 | 0.056 | 0.065 |
| A766 | 0.141 | 0.089 | 0.104 | 0.062 | N. A.* | 0.054 | 0.048 | 0.052 | 0.054 |
| A767 | 0.079 | 0.045 | 0.084 | 0.051 | 0.057 | 0.072 | 0.066 | 0.054 | 0.054 |
| **Cutoff value** (mean + 2*SD of healthy control) | | | | | | | | | |
| | 0.1122 | 0.1123 | 0.1073 | 0.0820 | 0.0867 | 0.0783 | 0.0784 | 0.0604 | 0.0916 |

* N. A., not available. Because these values are abnormally high (over 0.65), they were considered false values and not included in subsequent statistical analysis.

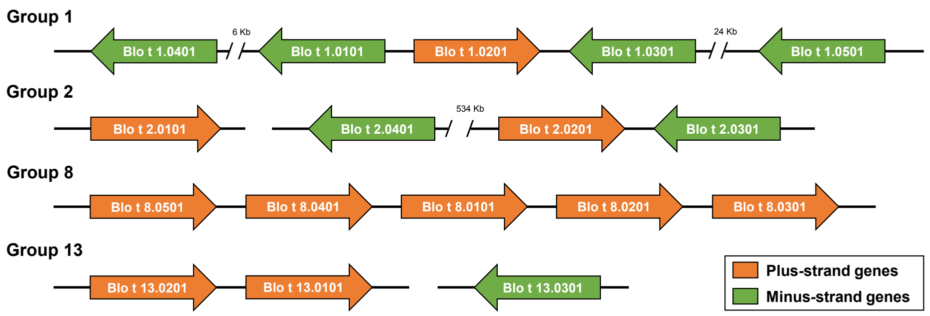


**Figure S1. Gene synteny of homologous allergen genes**

Gene synteny was aligned for the homologous genes of the group 1, 2, 8 and 13 allergens of *B. tropicalis*. All the gene IDs are consistent with those in Table 1. The gene distances without label are below 5 kb, which are regarded as tandem array.

**A**

```
CLUSTAL O(1.2.4) multiple sequence alignment


Blo_t_2.0101      MFKFICLALLPFWSRAAAGDVKFTDCAHGEVTSLDLSGCSGDHCIIHKGKSFTLKTFFIA  60
BT_003416.02      MFKFICLALLVSY--AAAGDVKFTDCAHGEVTSLDLSGCSGDHCTIHKGKSFTLKTFFIA  58
                  **********  :  ***************************** ***************

Blo_t_2.0101      NQDSEKLEIKISAIMNNIEVPVPGVDKDGCKHTTCPLKKGQKYELDYSLIIPTILPNLKT  120
BT_003416.02      NQDSEKLEIKISANMNGIEVPVPGVDKDGCKHTTCPLKKGQKYELDYSLIIPTFLPSVKT  118
                  ************* **.************************************:**.:**

Blo_t_2.0101      VTTASLVGDHGVVACGKVNTEVVD      144
BT_003416.02      VTTASLVGDHGVVACGKVNTEVVD      142
                  ************************
```

**B**

```
CLUSTAL O(1.2.4) multiple sequence alignment


Blo_t_2.0101      ATGTTCAAGTTTATCTGTCTCGCCCTTTTGCCCTTTTGGTCTCGTGCCGCCGCCGGTGAT  60
BT_003416.02      ATGTTCAAGTTTATCTGTCTCGCCCTTTT------GGTCTCGTACGCCGCCGCCGGTGAT  54
                  *****************************          **    ***************

Blo_t_2.0101      GTCAAATTTACCGATTGTGCACATGGTGAGGTTACCTCATTGGACTTGTCTGGATGCTCT  120
BT_003416.02      GTCAAATTTACCGATTGTGCACATGGTGAGGTTACCTCATTGGACTTGTCTGGATGCTCT  114
                  ************************************************************

Blo_t_2.0101      GGCGACCATTGCATAATCCACAAGGGAAAGAGCTTTACCTTGAAGACCTTTTTCATTGCT  180
BT_003416.02      GGCGACCATTGCACAATCCACAAGGGAAAGAGCTTTACCTTGAAGACCTTTTTCATTGCT  174
                  ************* **********************************************

Blo_t_2.0101      AACCAAGACTCTGAAAAGTTGGAGATCAAGATCAGCGCCATCATGAACAATATTGAGGTT  240
BT_003416.02      AACCAAGACTCTGAAAAGTTGGAGATCAAGATCAGCGCCAACATGAACGGTATTGAGGTT  234
                  **************************************** *******  **********

Blo_t_2.0101      CCAGTTCCAGGTGTCGACAAGGACGGTTGCAAGCACACCACCTGCCCATTGAAGAAGGGA  300
BT_003416.02      CCAGTTCCAGGTGTCGACAAGGACGGTTGCAAGCACACCACCTGCCCATTGAAGAAGGGA  294
                  ************************************************************

Blo_t_2.0101      CAAAAGTACGAACTCGACTACAGTTTGATCATCCCAACCATCTTGCCAAACTTGAAGACC  360
BT_003416.02      CAAAAGTACGAACTCGACTACAGTTTGATCATCCCAACCGTCTTGCCAAACGTGAAGACC  354
                  *************************************** *********** ********

Blo_t_2.0101      GTCACCACCGCATCGTTGGTTGGCGATCACGGTGTCGTTGCTTGCGGAAAGGTCAACACC  420
BT_003416.02      GTCACCACCGCATCGTTGGTTGGCGATCACGGTGTCGTTGCTTGCGGAAAGGTCAACACC  414
                  ************************************************************

Blo_t_2.0101      GAGGTTGTCGATTAA      435
BT_003416.02      GAGGTTGTCGATTAA      429
                  ***************
```

**Figure S2. Sequence alignment of the group 2 allergen of *B. tropicalis***
**(A)** Protein sequence alignment of the reported Blo t 2.0101 (GenBank accession: AAQ73483) and BT_003416.02 identified in the genome (Table 1). A mismatch region was marked in grey background. **(B)** Coding sequence alignment of the reported Blo t 2.0101 (GenBank accession: AY288141) and BT_003416.02 identified in the genome (Table 1). A mismatch region was marked in grey background and partial frame-shift matched sequences were linked by white lines.

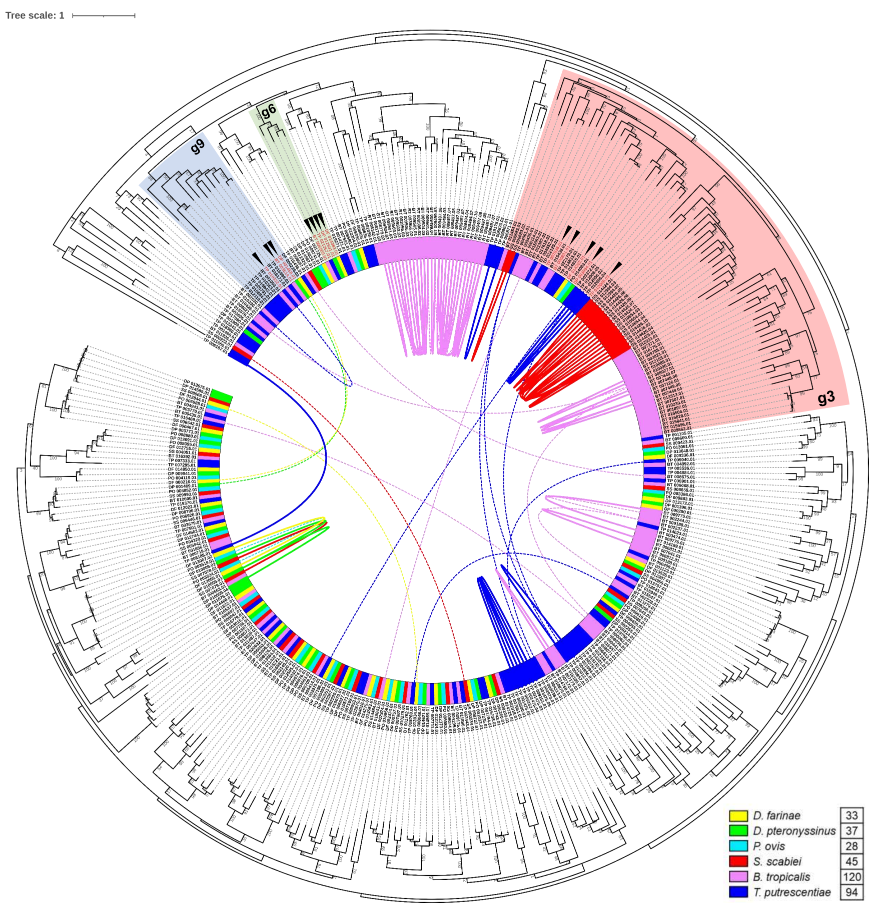


**Figure S3. Serine protease allergens in the phylogenetic tree**

The phylogenetic tree was adapted from Figure 4B in the previous report by Xiong Q. et al., 2022. Three clusters g3, g6 and g9 were highlighted that contain the group 3, 6 and 9 allergens, respectively. All the reported allergens were highlighted in red and marked with black triangles.

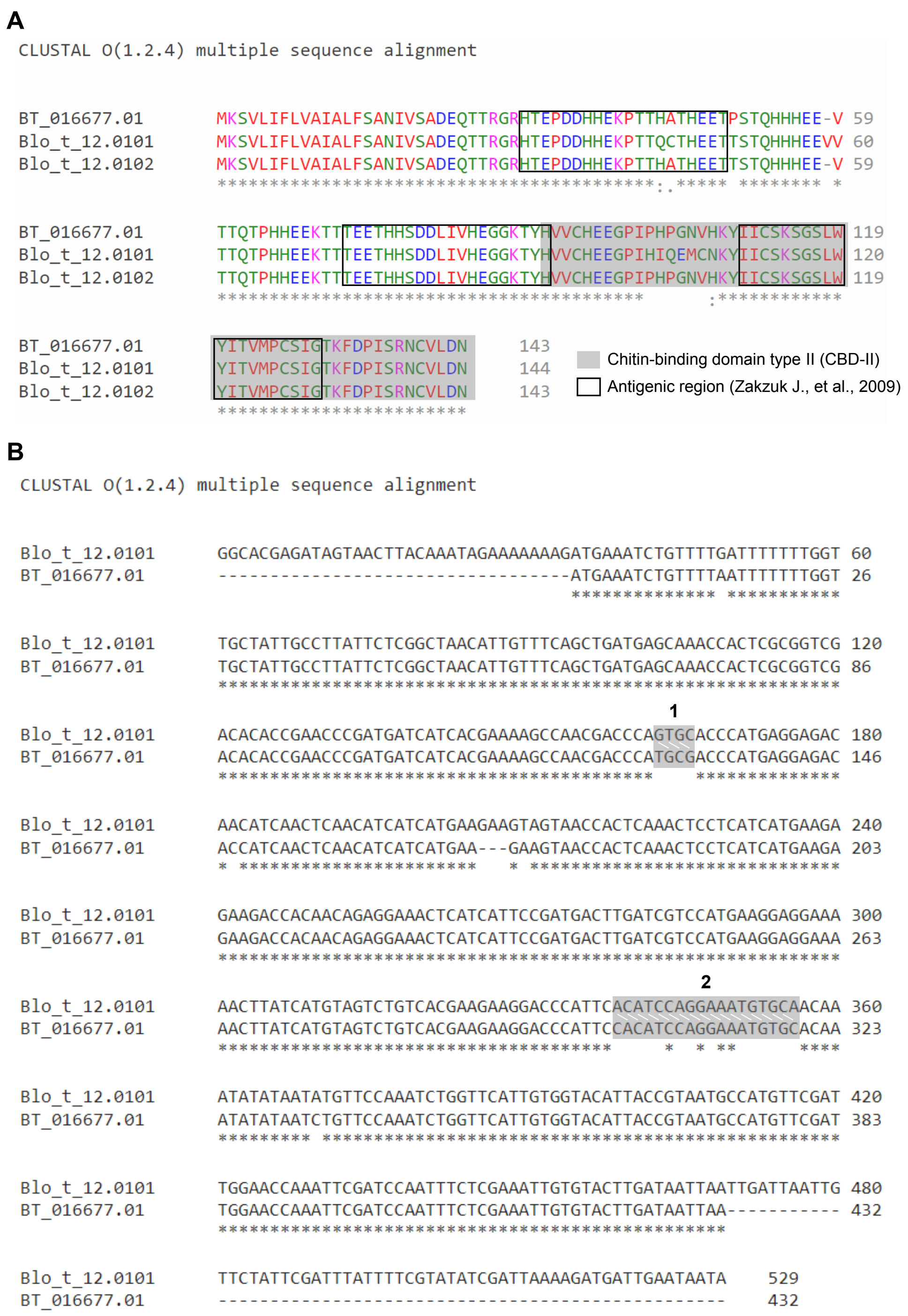


**Figure S4. Sequence alignment of the group 12 allergens of *B. tropicalis***
**(A)** Protein sequence alignment of BT_016677.01 identified in the genome (Table 1), the reported Blo t 12.0101 (GenBank accession: AAA78904) and Blo t 12.0102 reported by Zakzuk J. et al., 2009. Chitin-binding domain and antigenic regions were highlighted. **(B)** Coding sequence alignment of the reported Blo t 12.0101 (GenBank accession: U27479) and BT_016677.01 identified in the genome (Table 1). Two mismatch regions were marked in grey background and partial frame-shift matched sequences were linked by white lines.

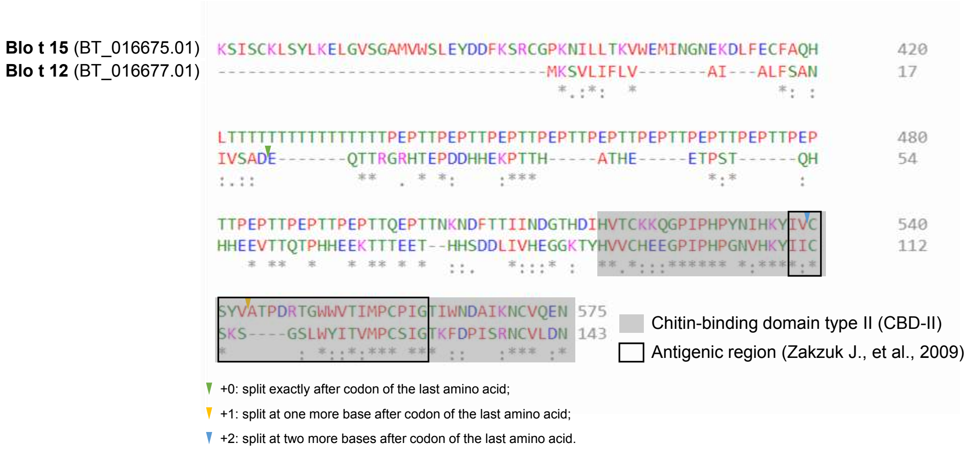


**Figure S5. Sequence alignment of Blo t 12 and 15**

Protein sequence alignment of Blo t 12 (gene locus: BT_016677.01) and Blo t 15 (gene locus: BT_016675.01) identified in the genome (Table 1). Chitin-binding domain and antigenic regions were highlighted. The splice sites (locations of exon–intron boundaries) were marked by colored triangles, but none of common splice sites was identified.

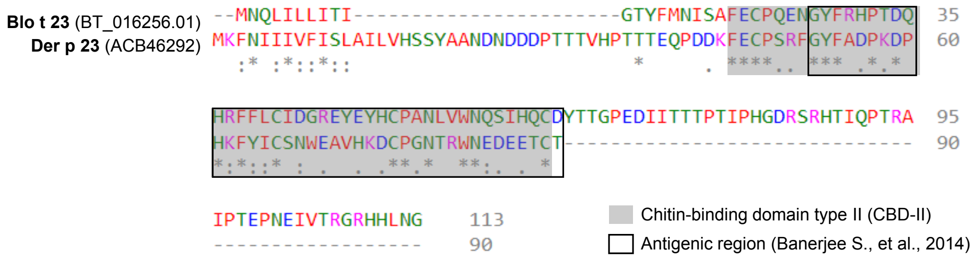


**Figure S6. Sequence alignment of Blo t 23 and Der p 23**
Protein sequence alignment of Blo t 23 (gene locus: BT_016256.01) identified in the genome (Table 1) and the reported Der p 23 (GenBank accession: ACB46292). Chitin-binding domain and antigenic regions were highlighted.

**A**

```
CLUSTAL O(1.2.4) multiple sequence alignment


DF_000078.02        --MKTIYAI-LSIMACIGLMNASIKRDHNDYSKNPMRIVCYVGTWSVYHKVDPYTIEDID  57
DF_000078.03        MKMKTTFAL-FCICFCIGLLNAATKRDHNDYSKNPMRIVCYVGTWSVYHKVDPYTIEDID  59
DP_005782.02        --MKTTLAL-FCIWACVGLMNAATKRDHNNYSKNPMRIVCYVGTWSVYHKVDPYTIEDID  57
DP_005782.03        --MKTSCAI-LILMACFGLMNAAVKRDHNNYSKNPMRIVCYVGTWSVYHKVDPYTIEDID  57
PO_009973.02        -MIKTKFTLFFLFFTIIGCINAVQKRDHNNYKKNPMRIVCYVGTWSVYHKVDPFQIEDID  59
BT_016675.01        --MRLLISI-VVLGTLLGSLTAVQKRDHNDYSKN-KRVVCYVGTWSVYHKNDPFEIEDID  56
TP_009563.02        --MRLLLSI-VLLGCLLGSLSAVQKRDRNNYTKNPKRVVCYLGTWSVYHKIDPFVIEDID  57
TP_009564.02        --MRLLLSI-VLLGCLLGSLSAVQKRDRNNYTKNPKRVVCYLGTWSVYHKIDPFVIEDID  57
                      ::   :: . :   .* :.*  ***:*:*.**  *:***:******** **: *****

DF_000078.02        PFKCTHLMYGFAKIDEYKYTIQVFDPYQDDNHNSWEKRGYERFNNLRLKNPELTTMISLG  117
DF_000078.03        PFKCTHLMYGFAKIDEYKYTIQVFDPYQDDNHNSWEKRGYERFNNLRLKNPELTTMISLG  119
DP_005782.02        PFKCTHLMYGFAKIDEYKYTIQVFDPFQDDNHNSWEKHGYERFNNLRLKNPELTTMISLG  117
DP_005782.03        PFKCTHLMYGFAKIDEYKYTIQVFDPYQDDNHNSWEKHGYERFNNLRSKNPELTTMISLG  117
PO_009973.02        PFKCTHLMYGFAKIDEYKYTIQVFDPYQDDNKNSWEKHGYERFNNLRLKNPELTTMISLG  119
BT_016675.01        VTKCTHVMYGFAKIDEYKYTIEVFDPYQDDSANPWDKRGYERFNNLRKQNPNLSTLISLG  116
TP_009563.02        VHRCTHVMYGFAKIDEYKYTIQVFDPYQDDSANPWDKKGYQRFNDLRKINPELTTMISLG  117
TP_009564.02        VHRCTHVNYGFAKIDEYKYTIQVFDPYQDDSANPWDKKGYQRFNDLRKINPELTTMISLG  117
                      :***: *************:****:***. * *:*:**:***:**  **:*:*:****

DF_000078.02        GWYEGSEKYSDMAANPTYRQQFIQSVLDFLQEYKFDGLDLDWEYPGSRLGNPKIDKQNYL  177
DF_000078.03        GWYEGSEKYSDMAANPAYRQQFIQSVLDFLQEYKFDGLDLDWEYPGSRLGNPKIDKQNYL  179
DP_005782.02        GWYEGSEKYSDMAANPTYRQQFVQSVLDFLQEYKFDGLDLDWEYPGSRLGNPKIDKQNYL  177
DP_005782.03        GWYEGSEKYSDMAANPTYRHQFVQSVLDFLQEYKFDGLDLDWEYPGSRLGNPKIDKQNYL  177
PO_009973.02        GWYEGSEKYSDMAANVMYRKQFVQSVLEFLEEYQFDGLDLDWEYPGSRLGDPKVDKQNYL  179
BT_016675.01        GWYEGSEKYSDMARNPTYRKNFVESVVAFLQKYDFDGLDLDWEYPGSRLGDPRVDKSNYL  176
TP_009563.02        GWYEGSEKYSDMARNPTYRKNFVDSVIAFLAKYDFDGLDLDWEFPGSRLGDPAVDATNFL  177
TP_009564.02        GWYEGSEKYSDMARNPTYRKNFVDSVIAFLAKYDFDGLDLDWEFPGSRLGDPAVDATNFL  177
                    ************* *  **::*::**: ** :*.*********:******:* :*  *:*

DF_000078.02        ALVRELKDAFEPHGYLLTAAVSPGKDKIDRAYDIKELNKLFDWMNVMTYDYHGGWENFY-  236
DF_000078.03        ALVRELKDAFEPHGYLLTAAVSPGKDKIDRAYDIKELNKLFDWMNVMTYDYHGGWENFY-  238
DP_005782.02        TLVRELKEAFEPFGYLLTAAVSPGKDKIDVAYELKELNQLFDWMNVMTYDYHGGWENVF-  236
DP_005782.03        TLVRELKEAFEPFGYLLTAAVSPGKDKIDVAYELKELNQLFDWMNVMTYDYHGGWENVF-  236
PO_009973.02        CLIRELKEAFEPYGYLLTAAVSPGKDKIDRAYDIPELNKLFDWFNVMTYDYHGGWEDFY-  238
BT_016675.01        ELVRELKAALDPHGYLLTAAMSPGQPTIEKAYHIKELNELLDFMNIMTYDYHGAWENVIG  236
TP_009563.02        ELVRELKTAFEPHGYLLTSAMSPGKDKIDLAYHIKELNELLDFMNIMTYDYHGGWEDFY-  236
TP_009564.02        ELVRELKTAFEPHGYLLTSAMSPGKDKIDLAYHIKELNELFDFMNIMTYDYHGGWEDFY-  236
                     *:**** *::*.*****:*:***: .*: **.: ***:*:*::*:*******.**:.

DF_000078.02        GHNAPLYKRPDET---DELHTYFNVNYTMHYYLNNGATRDKLVMGVPFYGPLK-IEANSN  292
DF_000078.03        GHNAPLYKRPDET---DELHTYFNVNYTMHYYLNNGATRDKLVMGVPFYGRAWSIEDRSK  295
DP_005782.02        GHNAPLYKRPDET---DELHTYFNVNYTMHYYLNNGATRDKLVMGVPFYGRAWSIEDRSK  293
DP_005782.03        GHNAPLYKRPDET---DELHTYFNVNYTMHYYLNNGATRDKLVMGVPFYGRAWSIEDRSK  293
PO_009973.02        GHNAPLYKRPDET---DDLHTFFTVNYTMHYYLSKGASRDKLVMGVPFYGRAWSIEDRSK  295
BT_016675.01        GHNAPLYPSPPTDKIDPVLNHIFNVNYTMHYYLEHGATKDKMVMGIPFYGRGWSLESRDL  296
TP_009563.02        GHNAPLYQRPDEH---DPLYTYFTVNYTMHYYLSHGATRDKLVMGVPFYGRAWSMEDPSK  293
TP_009564.02        GHNAPLYQRPDEH---DPLYTYFTVNYTMHYYLSHGATRDKLVMGVPFYGRAWSMEDPSK  293
                    *******  *        *   *.*********.:**::**:***:****    :*  .

DF_000078.02        L--EIQPKACRHPGFISGEEGVLSYIELCQLFQK--EEWHIQYDEYYNAPYGYNDKIWVG  348
DF_000078.03        LKLGDPAKGMSPPGFISGEEGVLSYIELCQLFQK--EEWHIQYDEYYNAPYGYNDKIWVG  353
DP_005782.02        VKLGDPAKGMSPPGFITGEEGVLSYIELCQLFQK--EEWHIQYDEYYNAPYGYNGKIWVG  351
DP_005782.03        VKLGDPAKGMSPPGFITGEEGVLSYIELCQLFQK--EEWHIQYDEYYNAPYGYNDKIWVG  351
PO_009973.02        VKIGDPAKGMSPAGFISGEQGVLAYIELCQLFKK--EKWTIHYDEYYNAPYAYNDKVWVG  353
BT_016675.01        VRFGDIVKGKSPPGFITGEEGVLGYNELCQMMQNDSQIWTTHYDQYFEAPYAYNGDIWIG  356
TP_009563.02        HALRDSAKGKSPPGFITGEEGVLGYNELCQMFRNASQTWITHYDEYYEAPYAYNGQLWVG  353
TP_009564.02        HALKDSAKGKSPPGFITGEEGVLGYNELCQMFRNASQTWHVHYDEYYEAPYAYNGQLWVG  353
                           *.    ***:**:***.* ****::::  : *  :**:*::***.**..:*:*

DF_000078.02        YDDLASISCKLAFLKELGVSGVMVWSLENDDFKGHCGPKNPLLNKVHNMINGDEKNSFEC  408
DF_000078.03        YDDLASISCKLAFLKELGVSGVMVWSLENDDFKGHCGPKNPLLNKVHNMINGDEKNSFEC  413
DP_005782.02        YDDLASISCKLAFLKELGVSGVMIWSLENDDFKGHCGPKYPLLNKVHNMINGDEKNSYEC  411
DP_005782.03        YDDLASISCKLAFLKELGVSGVMIWSLENDDFKGHCGPKYPLLNKVHNMINGDEKNSYEC  411
PO_009973.02        YDDLASLSCKLSFLKQLGVSGVMVWSVEDDDFKGHCGPKYPLLNKVYDMINGNDKHSYEC  413
BT_016675.01        YEDLKSISCKLSYLKELGVSGAMVWSLEYDDFKSRCGPKNILLTKVWEMINGNEKDLFEC  416
TP_009563.02        YDDLKSLSCKLAFLKSLDVSGAMVWSLENDDFKERCGPKNVLLTKVWDMINGAEKHSMEC  413
TP_009564.02        YDDLKSLSCKLAFLKSLDVSGAMVWSLENDDFKERCGPKNVLLTKVWDMINGAEKHSMEC  413
                    *:** *:****::**.*.***.*:**:* **** :****  **.** :**** :*.  **
```

**Figure S7. Sequence alignment of group 15 allergen homologs**

**(A)** Protein sequences (partial, without N-terminus) of all the homologs of the group 15 allergens (Cluster 10 in Figure 2B) were aligned and suggested their high conservation among six astigmatic mites.

**B**

```
CLUSTAL O(1.2.4) multiple sequence alignment


AAD52672.1      --MKTIYAILSIMACIGLMNASIKRDHNDYSKNPMRIVCYVGTWSVYHKVDPYTIEDIDP  58
DF_000078.02    --MKTIYAILSIMACIGLMNASIKRDHNDYSKNPMRIVCYVGTWSVYHKVDPYTIEDIDP  58
DF_000078.03    MKMKTTFALFCICFCIGLLNAATKRDHNDYSKNPMRIVCYVGTWSVYHKVDPYTIEDIDP  60
                  *** :*::.*  ****:**: *************************************

AAD52672.1      FKCTHLMYGFAKIDEYKYTIQVFDPYQDDNHNSWEKRGYERFNNLRLKNPELTTMISLGG  118
DF_000078.02    FKCTHLMYGFAKIDEYKYTIQVFDPYQDDNHNSWEKRGYERFNNLRLKNPELTTMISLGG  118
DF_000078.03    FKCTHLMYGFAKIDEYKYTIQVFDPYQDDNHNSWEKRGYERFNNLRLKNPELTTMISLGG  120
                ************************************************************

AAD52672.1      WYEGSEKYSDMAANPTYRQQFIQSVLDFLQEYKFDGLDLDWEYPGSRLGNPKIDKQNYLA  178
DF_000078.02    WYEGSEKYSDMAANPTYRQQFIQSVLDFLQEYKFDGLDLDWEYPGSRLGNPKIDKQNYLA  178
DF_000078.03    WYEGSEKYSDMAANPAYRQQFIQSVLDFLQEYKFDGLDLDWEYPGSRLGNPKIDKQNYLA  180
                ***************:********************************************

AAD52672.1      LVRELKDAFEPHGYLLTAAVSPGKDKIDRAYDIKELNKLFDWMNVMTYDYHGGWENFYGH  238
DF_000078.02    LVRELKDAFEPHGYLLTAAVSPGKDKIDRAYDIKELNKLFDWMNVMTYDYHGGWENFYGH  238
DF_000078.03    LVRELKDAFEPHGYLLTAAVSPGKDKIDRAYDIKELNKLFDWMNVMTYDYHGGWENFYGH  240
                ************************************************************

AAD52672.1      NAPLYKRPDETDELHTYFNVNYTMHYYLNNGATRDKLVMGVPFYGRAWSIEDRSKLKLGD  298
DF_000078.02    NAPLYKRPDETDELHTYFNVNYTMHYYLNNGATRDKLVMGVPFYGPLK-IEANSNL--EI  295
DF_000078.03    NAPLYKRPDETDELHTYFNVNYTMHYYLNNGATRDKLVMGVPFYGRAWSIEDRSKLKLGD  300
                *********************************************    **  .*:*

AAD52672.1      PAKGMSPPGFISGEEGVLSYIELCQLFQKEEWHIQYDEYYNAPYGYNDKIWVGYDDLASI  358
DF_000078.02    QPKACRHPGFISGEEGVLSYIELCQLFQKEEWHIQYDEYYNAPYGYNDKIWVGYDDLASI  355
DF_000078.03    PAKGMSPPGFISGEEGVLSYIELCQLFQKEEWHIQYDEYYNAPYGYNDKIWVGYDDLASI  360
                  *.   *****************************************************

AAD52672.1      SCKLAFLKELGVSGVMVWSLENDDFKGHCGPKNPLLNKVHNMINGDEKNSFECILGPSTT  418
DF_000078.02    SCKLAFLKELGVSGVMVWSLENDDFKGHCGPKNPLLNKVHNMINGDEKNSFECILGPSTT  415
DF_000078.03    SCKLAFLKELGVSGVMVWSLENDDFKGHCGPKNPLLNKVHNMINGDEKNSFECILGPSTT  420
                ************************************************************

AAD52672.1      TPTPTTTPTTPTTTPTTPSPTTPTTTPSPTTPTT---------TPSPTTPTTTPSPTTPT  469
DF_000078.02    TPTPTTTPTTPTTTPTTPSPTTPTTTPSPTTPTTTPSPTTPTTTPSPTTPTTTPSPTTPT  475
DF_000078.03    TPTPTTTPTTPTTTPTTPSPTTPTTTPSPTTPTTTPSPTTPTTTPS---------PTTPT  471
                **********************************         ***         *****

AAD52672.1      PTTPTPAPTTSTPSPTTTEHTSETPKYTTYVDGHLIKCYKEGDIPHPTNIHKYLVCEFVN  529
DF_000078.02    PTTTTPAPTTSTPSPTTTEHTSETPKYTTYVDGHLIKCYKEGDIPHPTNIHKYLVCEFVN  535
DF_000078.03    PTTPTPTPYNIDTFANHDRTHKRN----------------------TKIHHLYRWTFD-  507
                *** **:* .     .  .  ...                      *:**:     *

AAD52672.1      GGWWVHIMPCPPGTIWCQEKLTCIGE    555
DF_000078.02    GGWWVHIMPCPPGTIWCQEKLTCIGE    561
DF_000078.03    --------------------------    507
```

**Figure S7. Sequence alignment of group 15 allergen homologs**

**(B)** Protein sequences of all the homologs of the group 15 allergens of *D. farinae*. AAD52672.1 is the NCBI GenBank accession of the reported group 15 allergen of *D. farinae*, Der f 15.0101. The variable N-terminus in red square and higher similarity suggested DF_000078.02 to be the reported Der f 15, although DF_00078.03 was the one with higher similarity in another variable region in black squares. The highly variable proline (P), serine (S), and threonine (T)-rich region was marked in grey background.

**C**

```
CLUSTAL O(1.2.4) multiple sequence alignment


AAY84565.1      MKTSCAILILMACFGLMNAAVKRDHNNYSKNPMRIVCYVGTWSVYHKVDPYTIEDIDPFK  60
AAY84564.2      MKTSCAILILMACFGLMNAAVKRDHNNYSKNPMRIVCYVGTWSVYHKVDPYTIEDIDPFK  60
DP_005782.02    MKTTLALFCIWACVGLMNAATKRDHNNYSKNPMRIVCYVGTWSVYHKVDPYTIEDIDPFK  60
DP_005782.03    MKTSCAILILMACFGLMNAAVKRDHNNYSKNPMRIVCYVGTWSVYHKVDPYTIEDIDPFK  60
                ***: *:: : **.******.**************************************

AAY84565.1      CTHLMYGFAKIDEYKYTIQVFDPYQDDNHNTWEKHGYERFNNLRLKNPELTTMISLGGWY  120
AAY84564.2      CTHLMYGFAKIDEYKYTIQVFDPYQDDNHNTWEKHGYERFNNLRLKNPELTTMISLGGWY  120
DP_005782.02    CTHLMYGFAKIDEYKYTIQVFDPFQDDNHNSWEKHGYERFNNLRLKNPELTTMISLGGWY  120
DP_005782.03    CTHLMYGFAKIDEYKYTIQVFDPYQDDNHNSWEKHGYERFNNLRSKNPELTTMISLGGWY  120
                ***********************:******:************* ***************

AAY84565.1      EGSEKYSDMVANPTYRKQFVQSVLDFLQEYKFDGLDLDWEYPGSRLGNPKIDKQNYLTLV  180
AAY84564.2      EGSEKYSDMVANPTYRQQFVQSVLDFLQEYKFDGLDLDWEYPGSRLGNPKIDKQNYLTLV  180
DP_005782.02    EGSEKYSDMAANPTYRQQFVQSVLDFLQEYKFDGLDLDWEYPGSRLGNPKIDKQNYLTLV  180
DP_005782.03    EGSEKYSDMAANPTYRHQFVQSVLDFLQEYKFDGLDLDWEYPGSRLGNPKIDKQNYLTLV  180
                *********.******:*******************************************

AAY84565.1      RELKEAFEPFGYLLTAAVSPGKDKIDVAYELKELNQLFDWMNVMTYDYHGGWENVFGHNA  240
AAY84564.2      RELKEAFEPFGYLLTAAVSPGKDKIDVAYELKELNQLFDWMNVMTYDYHGGWENVFGHNA  240
DP_005782.02    RELKEAFEPFGYLLTAAVSPGKDKIDVAYELKELNQLFDWMNVMTYDYHGGWENVFGHNA  240
DP_005782.03    RELKEAFEPFGYLLTAAVSPGKDKIDVAYELKELNQLFDWMNVMTYDYHGGWENVFGHNA  240
                ************************************************************

AAY84565.1      PLYKRPDETDELHTYFNVNYTMHYYLNNGATRDKLVMGVPFYGRAWSIEDRSKVKLGDPA  300
AAY84564.2      PLYKRPDETDELHTYFNVNYTMHYYLNNGATRDKLVMGVPFYGRAWSIEDRSKVKLGDPA  300
DP_005782.02    PLYKRPDETDELHTYFNVNYTMHYYLNNGATRDKLVMGVPFYGRAWSIEDRSKVKLGDPA  300
DP_005782.03    PLYKRPDETDELHTYFNVNYTMHYYLNNGATRDKLVMGVPFYGRAWSIEDRSKVKLGDPA  300
                ************************************************************

AAY84565.1      KGMSPPGFITGEEGVLSYIELCQLFQKEEWHIQYDEYYNAPYGYNDKIWVGYDDLASISC  360
AAY84564.2      KGMSPPGFITGEEGVLSYIELCQLFQKEEWHIQYDEYYNAPYGYNDKIWVGYDDLASISC  360
DP_005782.02    KGMSPPGFITGEEGVLSYIELCQLFQKEEWHIQYDEYYNAPYGYNGKIWVGYDDLASISC  360
DP_005782.03    KGMSPPGFITGEEGVLSYIELCQLFQKEEWHIQYDEYYNAPYGYNDKIWVGYDDLASISC  360
                *********************************************.**************

AAY84565.1      KLAFLKELGVSGVMIWSLENDDFKGHCGPKYPLLNKVHNMINGDEKNSYECLLGPSTTTP  420
AAY84564.2      KLAFLKELGVSGVMIWSLENDDFKGHCGPKYPLLNKVHNMINGDEKNSYECLLGPSTTTP  420
DP_005782.02    KLAFLKELGVSGVMIWSLENDDFKGHCGPKYPLLNKVHNMINGDEKNSYECLLGPSTTTP  420
DP_005782.03    KLAFLKELGVSGVMIWSLENDDFKGHCGPKYPLLNKVHNMINGDEKNSYECLLGPSTTTP  420
                ************************************************************

AAY84565.1      TPTTPSTPSTTTPTPTTPSTPSTT--------------------------TPTPTTPSTP  454
AAY84564.2      TPTTPSTPSTTTPTPTTPSTPSTTTPTPTTPSTPSTTTPTPTTPSTPSTTTPTPTTPSTT  480
DP_005782.02    TPTTPSTPSTTTPTPTTPSTPSTTTPTPTTPSTPSTTTPTPTTPSTPSTTTPTPTTPSTP  480
DP_005782.03    TPTTPSTPSTTTPTPTTPSTPSTTTPTP-------------TTPSTPSTTTPTPTTPSTT  467
                ************************                          ********* 

AAY84565.1      STTTPTPTTTDSTSETPKYTTYVDGHLIKCYKEGDLPHPTNIHKYLVCEYV---NGGWWV  511
AAY84564.2      STTTPTPTTTDSTSETPKYTTYVDGHLIKCYKEGDLPHPTNIHKYLVCEYV---NGGWWV  537
DP_005782.02    STTTPTPTTTDSTSETPKYTTYIDGHLIKCYKQGYLPHPTDVHKYLVCEYIATPNGGWWV  540
DP_005782.03    STTTPTPTTTDSTSETPKYTTYVDGHLIKCYKEGDLPHPTNIHKYLVCEYV---NGGWWV  524
                **********************:*********:* *****::********:   ******

AAY84565.1      HIMPCPPGTIWCQEKLTCITE 532
AAY84564.2      HIMPCPPGTIWCQEKLTCITE 558
DP_005782.02    HIMDCPKGTRWHATLKNCIQE 561
DP_005782.03    HIMPCPPGTIWCQEKLTCITE 545
                *** ** ** *     .** *
```

**Figure S7. Sequence alignment of group 15 allergen homologs**

**(C)** Protein sequences of all the homologs of the group 15 allergens of *D. pteronyssinus*. AAY84565.1 and AAY84564.2 are the NCBI GenBank accessions of the two reported group 15 isoallergens of *D. pteronyssinus*, Der p 15.0101 and Der p 15.0102, respectively. The two variable regions in red squares suggested DP_005782.03 to be the reported Der p 15. The highly variable P, S, and T-rich region was marked in grey background.

A

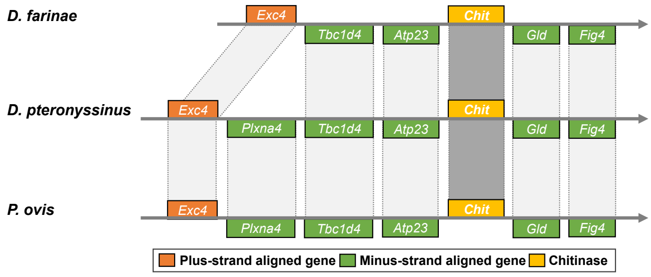


B

```
CLUSTAL O(1.2.4) multiple sequence alignment


DF_004204.02        --MTRFSLTVLAVLAACF-GSNIRPNVATLEPKTVCYYESWVHWRQGEGKMDPEDIDTSL   57
DP_004892.02        --MTRLSFTVLIFLAAYF-GSNIRPNVATLDPKTVCYYESWVHWRQGDGKMDPEDIDTSL   57
PO_003203.01        --MAKLSLQIIILLVIYSCSYLIEPNSATENPKTVCYYESWVYWRKGEGKMEPKDIDTSL   58
BT_010400.01        --MTKLCLVALATLAT--IAALVPVNHATKDPKTVCYYESWVHWRHGDGKMEPNEIDTSL   56
TP_022773.01        DQSSKLSLVALVATVL-AATLNIEPVQSTSSPKTVCYYEYWVHWRTGDGKQDPSEIDYTL   59
TP_001639.01        --MAKLCLLALASLAL--VVALMPSSEATSHPKTVCYYESWVHWRHGDGHMEPNEIDTTL   56
TP_021623.01        --MAKLCLLALASLAL--VVALMPSSEATSHPKTVCYYESWVHWRHGDGHMEPNEIDTTL   56
                       :::.:  :   .       :    :*  ******** **:** *:*: :*.:** :*

DF_004204.02        CTHIVYSYFGIDAATHEIKLLDEYLMKDLHDMEHFTQHKGNAKAMIAVGGSTMSDQFSKT   117
DP_004892.02        CSHIVYSYFGIDASSHEIKLLDQYLMITLHDMEHFTKHKGNAKAMIAVGGASMSDQFSKT   117
PO_003203.01        CTHIVYSYFGIDENTHEIKLLDPYLMLDLHEMEHFTKHKGNAKALIAIGGSTMSNQFSKT   118
BT_010400.01        CTHIVYTYFGIDAITHELKWLDPYLMKDLHDIEKFVQAKGKAKAMIAIGGASMSDQFSIT   116
TP_022773.01        CTHIVYSYMGIDGGTHEIKLLDPYLMNDLHDIDKFAKAKGNSKALVSVGGAAASSQFSIT   119
TP_001639.01        CTHIVYAYFGIDASTHELKWLDPYLSKDLHDLEKFVKAKGSAKALIAIGGASMSEQFGVT   116
TP_021623.01        CTHIVYAYFGIDASTHELKWLDPYLSKDLHDLEKFVKAKGSAKALIAIGGASMSEQFGVT   116
                    *:****:*:***  :**:* ** **   **::::*.: **.:**::::**:: *.**. *

DF_004204.02        AAVEHYRETFVVSTVDLMTRYGFDGVMIDWSGMQAKDSDNFIKLLDKFDEKFAHTSFVMG   177
DP_004892.02        AAVEHYRETFVVSTIDLMTKYGFDGVMIDWSGMQAKDSDNFVKLLDKFDEKFAQTSFVMG   177
PO_003203.01        AAVEHYREIFAASTIDLMTRYNFDGLMIDWSNEEAKDSDNLLKLLDKFDEKLDSTSFIMG   178
BT_010400.01        AGNEQYRDIFARSVVNFLAQYHFDGIMIDWYGVQERDSENLIHLLDKFDEKFASTTYSMG   176
TP_022773.01        VSVDSYRTAFVNSVVSFISSHNFDGVILDWSGMTASDNAHFIALLDKFDEKLANTPYTLA   179
TP_001639.01        AGSDQYREAIARSVVSFCAQYQFDGVMIDWSGIAEKDSDNLIKLLDKFDEKFASTPYTLG   176
TP_021623.01        AGSDQYREAIARSVVSFCAQYQFDGVMIDWSGIAEKDSDNLIKLLDKFDEKFASTPYTLG   176
                    .. : **  :. *.:.: : : ***:::** .     *. ::: ********:  * : :.

DF_004204.02        VTLPATIASYDNYNIPAISNYVDFMNVLSLDYTGSWAHTVGHASPFPEQLKTLEAYHKRG   237
DP_004892.02        VTLPATIASYDNYNIPAISNYVDFMNVLTLDYDGPWAYTVGHASALPEQLKTLEAYNKRG   237
PO_003203.01        ISLPATIASYDNYDIPKINDYVDLFNIITLDYTGSWAHTVGHASPFPDQLKTLEAYNKRG   238
BT_010400.01        ITLPATVATLDHYNVPKITVYVDFINVLTLDYAGPWGKVVDNASPLPEQLKTMEEYHKRG   236
TP_022773.01        VTVPASLNT-AAYNVPKIAQYVDFINVIAHDYAGPWSKQVGHASLLSHALSTMEAYHKAG   238
TP_001639.01        ITLPASTVTIQAVNVPKVTQYVDFINVLTIDYAGPWAKVVGHASPLPEQLKTLEEYHKRG   236
TP_021623.01        ITLPASTVTIQAVNVPKVTQYVDFINVLTIDYAGPWAKVVGHASPLPEQLKTLEEYHKRG   236
                    :::**:  :    ::* :  ***::*::: ** * *.  *.:** : . *.*:* *:* *

DF_004204.02        APRHKMVMAVPFYARTWILEKMNKQDIGDKASGPGPRGQFTQTDGFLSYNELCVQIQAET   297
DP_004892.02        APRHKMVMAVPFFARTWILEKMDKQDVGDKASGPGPKGQFTQTPGFLSYNELCVQIQAET   297
PO_003203.01        APRHKMVMAVPFYARTWMLAKINKQDVGDRAIGPGPKGQYTQTDGLLSYNELCVQILSEQ   298
BT_010400.01        APRSKLVMAVPMYARTWRLASPLHQDLGDAAISGGTKGPYTDTEGILSYNELCVRIKGSP   296
TP_022773.01        APRSKLVMSVPAFARTWRLANPAKQDIGDEALGGGEKGPYTQVEGLLSYNELCVLMRAHV   298
TP_001639.01        APRSKLVMAVPLYARTWKLASPAHQDIGDAAVSGGAKGPYTDVEGLLSYNELCVQIKQNP   296
TP_021623.01        APRSKLVMAVPLYARTWKLASPAHQDIGDAAVSGGAKGPYTDVEGLLSYNELCVQIKQNP   296
                    *** *:**:** :**** *  .  :**:** *  . * :* :*:. *:******** :

DF_004204.02        NAFTITRDHDNTAIYAVYVHS-NHAEWISFEDRHTLGEKAKNITQQGYAGMSVYTLSNED   356
DP_004892.02        NAFSITRDHDNTAIYAVYVHD-NHAEWISFEDRHTLGDKARNITEQGYGGMSVYTLSNED   356
PO_003203.01        SAFSIVRDEANTAIYAVYVHG-NEAEWISFEDRKTLAIKARNITQLGYAGMTVYTLSNED   357
BT_010400.01        NSFNIVRDVANTAVHAVYLHG-NEAEFYSFEDTKTLAAKAHNVTTMGYGGLSIFTLSNED   355
TP_022773.01        SQFSLVRDPAEHAVYAVYLRDSGHAEWLSFEDSKSLAFKADNVTRSGYAGLSVYSISNED   358
TP_001639.01        AAYSIIRDHANTAVHAVHLSG-STAEFISFEDSKTLAEKAKNITTEGYAGISVYTLSNED   355
TP_021623.01        AAYSIIRDHANTAVHAVHLSG-STAEFISFEDSKTLAEKAKNITTEGYAGISVYTLSNED   355
                      :.: **  : *::**:: . . **: **** ::*. ** *:*  **.*::::::****

DF_004204.02        VHGVCGDKNPLLHAIQSNYYHGVVTEPTVVTL-PPVTHTTEHVTDIPGVFHCHEEGFFRD   415
DP_004892.02        VHGVCGDKNPLLHAINSNYFRGIVTEPTVVTV-TPVTHTTEHVTDIPGVFHCHQEGFFRD   415
PO_003203.01        VHGVCGNKDPLLHAINENYYRGVVTEPPISTL-PPVTHTTEHVTDIPGVFHCHAEGFFRD   416
BT_010400.01        SHGTCGKKYPLLHSIVENYNHDPIDEVPITTIPPPTPHPTEVVTDIPGVFKCHSVGKFRD   415
TP_022773.01        VHGVCGQKNPLLHAINAHYWRGAVTEGPT-----PPNHFTPGPT--------EAPGKFRD   405
TP_001639.01        VHGTCGKKYPLVHAIVENYNKADVHEVPITTLSPPVTHPATPPPTVPGVFKCDHEGKFRD   415
TP_021623.01        VHGTCGKKYPLVHAIVENYNKADVHEVPITTLSPPVTHPATPPPTVPGVFKCDHEGKFRD   415
                     **.**.* **:*:*  :* :  : *        *  * :              .  * ***

DF_004204.02        KTYCATYYECKKGDFG-LEKTVHHCANHLQAFDEVSRTCIDHTKIPGC--   462
DP_004892.02        KTYCAKYYECKKGDFG-LEQTVHHCPNHSQAFDEVSRTCVDHTKIPGC--   462
PO_003203.01        KIYCSKYYECKKSDFG-LEMTVHQCPDHKQAFDEESKSCVDHTKVPGC--   463
BT_010400.01        HEYCFKYYDCVMGDFG-LESTVMYCERH-QAFDEKTYKCVEASQIPGC--   461
TP_022773.01        PVYCFKYYDCEKGEFGGFESTVFYCERH-QAWDEHAQKCVDAKTVPGC--   452
TP_001639.01        HQYCFKYYECVKGEFGGFEQTVMNCERH-QAFDEHLQKCVEAKTVPGCGN   464
TP_021623.01        HQYCFKYYECVKGEFGGFEQTVMNCERH-QAFDEHLQKCVEAKTVPGCGN   464
                      ** .**:*  .:** :* **  *  * **:**   .*:: . :***
```

**Figure S8. Sequence alignment of group 18 allergen homologs**

**(A)** Gene synteny alignment of the group 18 allergen loci of three astigmatic mites. None of conserved gene synteny was found in other astigmatic mites. Group 18 allergens are the chitinases. **(B)** Protein sequences of all the homologs of the group 18 allergens (Cluster 1 in Figure 2B) were aligned and suggested their high conservation among six astigmatic mites. Except DF_004204.02 and DP_004892.02 are reported Der f 18 and Der p 18, respectively, others are all putative allergens.

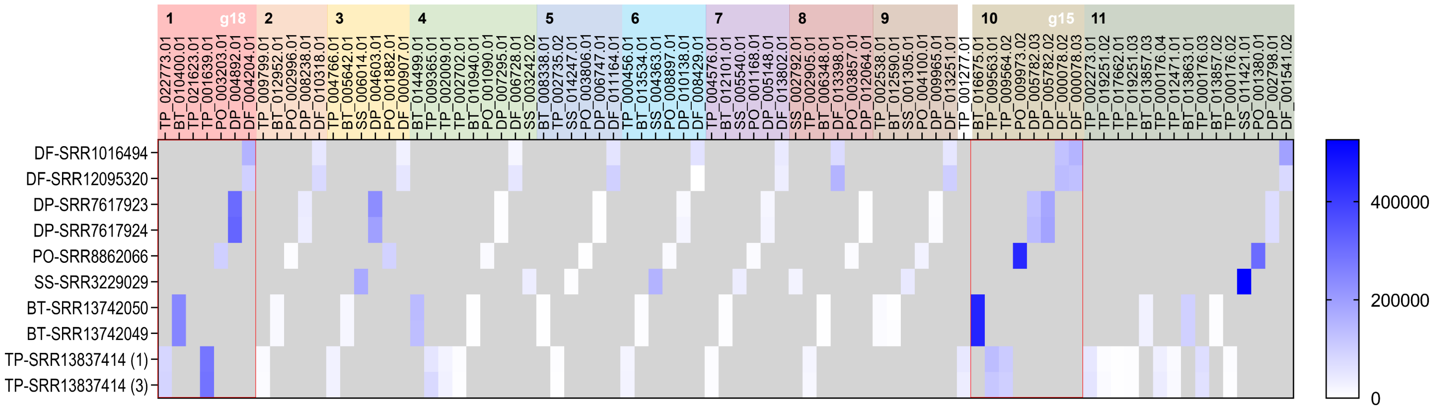


**Figure S9. Gene expression level of chitinases of six astigmatic mites**

Gene expression levels of chitinases of six astigmatic mites (Figure 2B) were quantified as TPM using transcriptome data of adult mites. The SRA accessions were labeled in the axis. Two clusters containing the group 15 and 18 allergens were highlighted in red squares. Two RNA transcriptome data of *B. tropicalis* (prefix: BT) were used, but not other two cDNA transcriptome data. TP1 and TP3 of SRR13837414 transcriptome data were used for genes of *T. putrescentiae* (labeled 1 and 3, respectively).

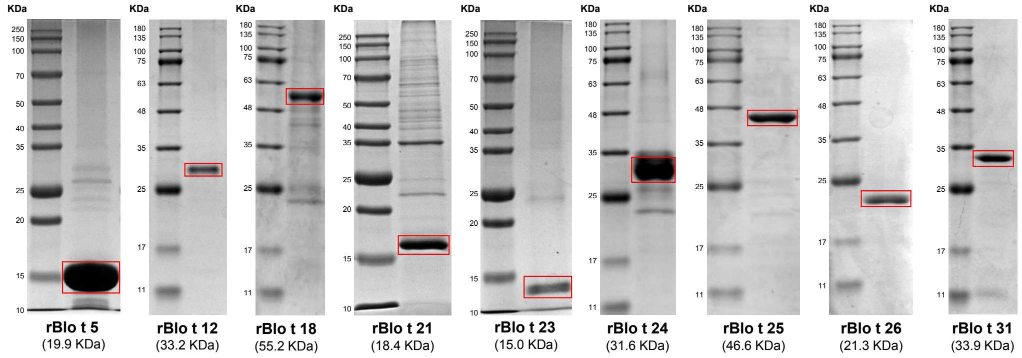


**Figure S10. SDS-PAGE analysis of nine purified recombinant proteins**

Nine purified recombinant proteins were loaded and analyzed in SDS-PAGE. All the nine recombinant proteins were observed within the range of estimated molecular weights (in the brackets under protein IDs) ± 5 KDa. The target bands of recombinant proteins were marked in red squares.

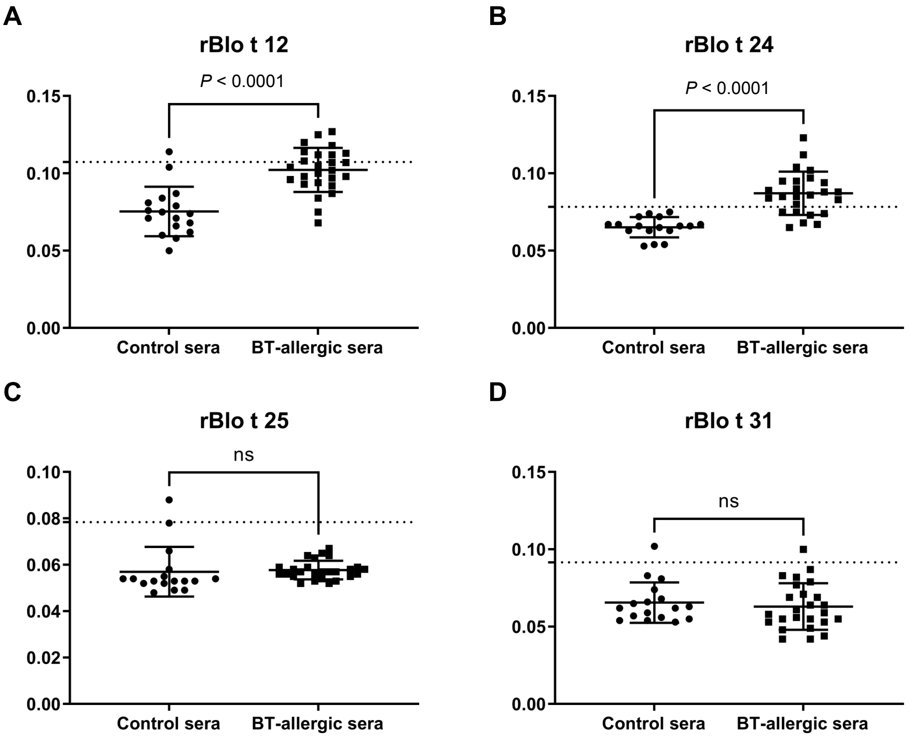


**Figure S11. ELISA results of four recombinant proteins for Blo t 12 and putative Blo t 24, 25 and 31**
The IgE antibody binding to recombinant proteins **(A)** rBlo t 12, **(B)** rBlo t 24, **(C)** rBlo t 25 and **(D)** rBlo t 31 was evaluated by ELISA using serum samples from 26 patients positive to *B. tropicalis* (BT-allergic sera) and 17 healthy individuals as control sera (Table S1). The absorbance values at 450 nm of recombinant proteins were recorded to represent the IgE levels of the putative allergens. The lines in dot plots were set as the mean with a standard deviation (SD) and the dotted lines indicated the cutoff value, mean + 2*SD of the healthy controls. Unpaired two-tailed *t*-test *p*-values indicate statistical significance (ns, not significant, $p > 0.5$). The recombinant proteins, rBlo t 12 and 24, presented significantly higher IgE levels in BT-allergic sera, but the low absorbance values were all below 0.15, which could not well confirm their allergenicity. Even though rBlo t 12, 24, 25 and 31 were expressed in pET-32a+ vector with a N-terminal thioredoxin tag that may cause false-positive IgE binding, we could not observe their significant allergenicity with these ELISA results.

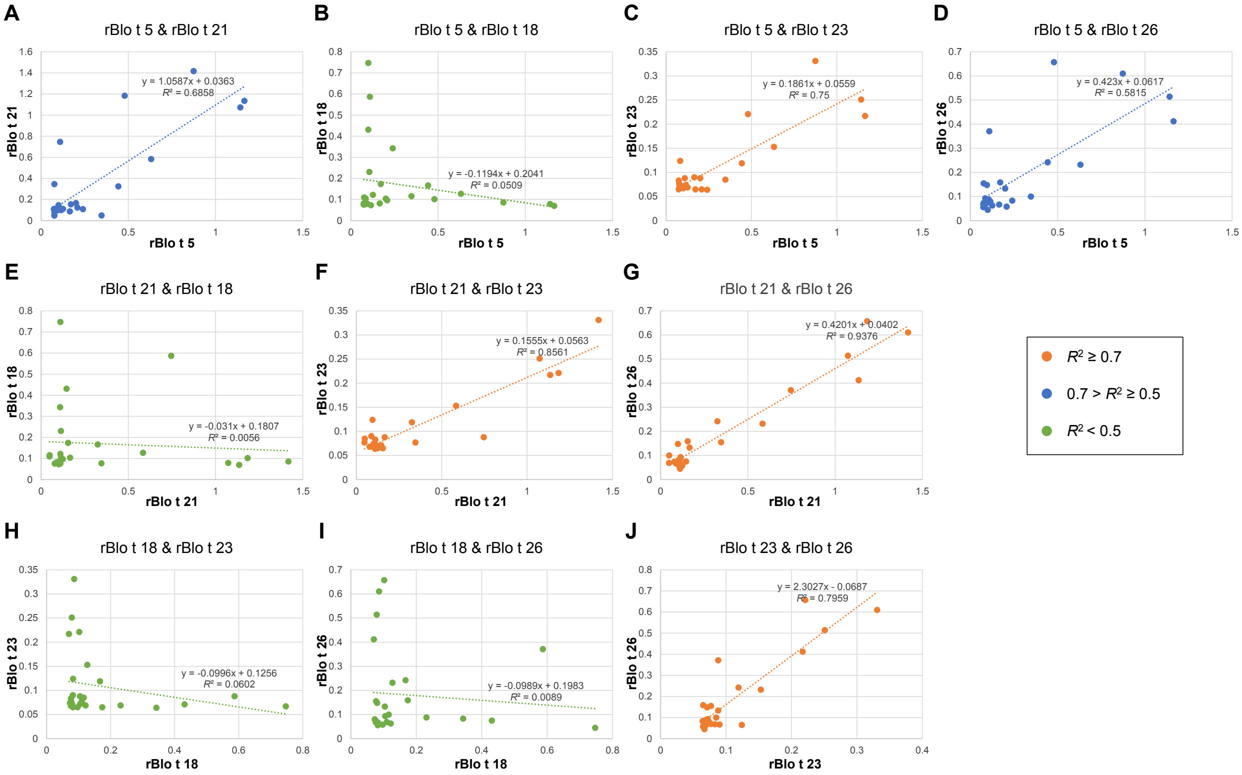


**Figure S12. Regression analysis of the IgE levels of five recombinant allergen proteins**
The IgE levels of five recombinant allergen proteins evaluated by ELISA (Figure 3) were subject to the regression analysis of ten pairs as follows: **(A)** rBlo t 5 & rBlo t 21, **(B)** rBlo t 5 & rBlo t 18, **(C)** rBlo t 5 & rBlo t 23, **(D)** rBlo t 5 & rBlo t 26, **(E)** rBlo t 21 & rBlo t 18, **(F)** rBlo t 21 & rBlo t 23, **(G)** rBlo t 21 & rBlo t 26, **(H)** rBlo t 18 & rBlo t 23, **(I)** rBlo t 18 & rBlo t 26 and **(J)** rBlo t 23 & rBlo t 26. $R^2$ values were used to assess correlation levels between a pair of recombinant proteins.
